# An integrated *in silico*-*in vitro* approach for identification of therapeutic drug targets for osteoarthritis

**DOI:** 10.1101/2021.09.27.461207

**Authors:** Raphaëlle Lesage, Mauricio N. Ferrao Blanco, Roberto Narcisi, Tim Welting, Gerjo J.V.M. van Osch, Liesbet Geris

## Abstract

Without the availability of disease-modifying drugs, there is an unmet therapeutic need for osteoarthritic patients. During osteoarthritis, the homeostasis of articular chondrocytes is dysregulated and a phenotypical transition called hypertrophy occurs, leading to cartilage degeneration. Targeting this phenotypic transition has emerged as a potential therapeutic strategy. Chondrocyte phenotype maintenance and switch are controlled by an intricate network of intracellular factors, each influenced by a myriad of feedback mechanisms, making it challenging to intuitively predict treatment outcomes. In this study, we developed a regulatory network model using knowledge-based and data-driven modelling technologies. The *in silico* high-throughput screening of (pairwise) perturbations operated with that network model highlighted conditions impacting the hypertrophic switch. Several combinations were tested in a murine cell line and primary chondrocytes to validate the predicted conditions’ potential. Our *in silico-in vitro* strategy opens a new route for developing osteoarthritis targeting therapies by refining the early stages of drug discovery.

## 1. Introduction

Osteoarthritis (OA) is a degenerative disease of the joint increasingly prevalent due to the ageing population. It is a major societal burden as no disease-modifying drugs are currently available on the market (*1*). OA is characterized by cartilage damage, led by an overall increase of catabolic processes and disturbance of anabolic processes. The joint cartilage is composed of a unique cell type, the chondrocyte, which is responsible for maintaining the tissue homeostasis in an environment mainly composed of water and biomolecules such as proteoglycans and collagen fibers. Many factors, including inflammation, may influence the shift from stable healthy cartilage towards a diseased state (*2*). Regardless of the exact inducing mechanisms, during that transition, some of the chondrocytes enter a maturation process called hypertrophy (*3*, *4*), leading to extracellular matrix (ECM) degradation, mineralization and bone formation. This pathological phenomenon resembles the hypertrophic changes observed in the course of endochondral ossification during growth and development (*2, 5*–*8*). Therefore, controlling the chondrocyte phenotype to prevent hypertrophic maturation has emerged as a potential therapeutic strategy to treat OA patients (*7*, *9*).

Crucial in this approach is the understanding of the process of articular chondrocyte hypertrophy for the identification of key regulators as potential drug targets. Several kfactors have been associated to the promotion of this phenotypic shift, such as Indian hedgehog (IHH) and inflammatory signalling pathways (*10*) and routes downstream of various growth factors are thought to be important in the control or disruption of chondrocyte homeostasis, such as the WNT and Bone morphogenic protein (BMP) pathways, the parathyroid hormone related peptide (PTHrP), as well as the insulin-like growth factor (IGF)-I, fibroblast growth factors (FGF) and transforming growth factors (TGF)-β (*9*, *11*, *12*). However, the interplay of intracellular pathways is highly intricate with extensive feedback loops, non-linear pathways, redundancy and intertwinings (*11*, *13*, *14*). This complicates the intuitive prediction of what will happen in case of perturbations of a specific target. For example, it was observed that the *in vitro* activation of the WNT pathway with the WNT3A ligand and the inhibition of that same pathway with Dickkopf1 (DKK1), both induced a reduction of glycosaminoglycan rich ECM in human articular chondrocytes (*15*). The fact that an activator and an inhibitor of the canonical WNT pathway both lead to the same outcome is surprising and highlights the intricacy of the underlying mechanisms. Hence, the ability to predict the effect of an external perturbation and potential therapies requires a systemic view on the process (*13*, *14*).

We propose to unravel the complexity of these regulatory pathways and to rationalize the identification of potential drug targets via screening of (combination) therapies by using a classical engineering approach, namely that of computer modeling and simulation. Contrary to *in vitro* and *in vivo* approaches, having a systemic view of the process using an *in silico* model allows to study the system numerically, in a cost and time efficient way and with less ethical concerns. In addition, it allows to prioritize experiments, thereby refining the traditional funnel of drug targets identification in the drug discovery process. The *in silico* approach starts with acquiring a systems perspective of the intracellular biological processes. It is necessary to identify the important individual components of the system and to know how they interact and influence each other. A computational model built on this information should generate results consistent with current knowledge but also allow to ask questions that would lead to new insights into yet unexplored situations and interactions (*16*). Such computational mechanistic approaches were already used in the past to identify influential candidates for cancer therapeutics (*17*), study the control stem cell fate decision (*18*) or prioritize personalized combination therapies (*19*).

In this study, we developed an *in silico* model of the regulatory network capturing articular chondrocyte phenotypic changes during OA. This network is built by combining knowledge-based modeling and data-driven approaches to ensure the mechanistic accuracy of the network whilst taking advantage of current automatic network reconstruction technologies. We characterized the numerical intracellular states of the articular chondrocyte model and looked into consistency with known physiological behaviors, through mathematical model implementation and computer simulations. We subsequently used the model as a ‘virtual chondrocyte’ to perform an *in silico* high throughput screening for early predictions of the best potential therapeutic conditions to be tested in wet lab experiments. Finally, we confirmed *in vitro* the role of predicted factors in the regulation of the phenotypic change, for both single and combinations of factors.

## 2. Results

### Highly interconnected intracellular networks regulate articular chondrocytes: building a mechanistic model

In order to build an articular chondrocyte model, we gathered known biological mechanisms from the literature into an activity flow graph or network (**Fig. 1**). In this graphical influence network, or activity flow representation, the nodes represent biological components (proteins or genes, see **Table S1**) important for chondrocyte biology (*9*). Directed edges linking two nodes represent interactions or activating/inhibitory influences between source and target proteins or between transcription factor (TFs) and target genes. Information was collected and adapted from a previously published model of growth plate chondrocytes (*20*, *21*) as well as from additional deep literature and database curation (see Method section). The annotations, descriptions and cross-references for each network node and its interactions can be consulted in two interactive networks, a protein signaling one and a gene regulatory one. The former goes from the growth factors binding their respective receptors down to the TF entering the nucleus. The latter represents a network of transcription factors regulating the expression of their target genes, coding for the corresponding proteins in the signaling network (**Fig. 1**). The two networks are interconnected as each biological component is represented by a gene in the gene regulatory network (GRN) and its corresponding protein in the signaling network. The combination of both networks can be regarded as an interactive knowledge base on chondrocyte signaling and is available through the online platform Cell Collective (*22*). We refer the reader to that interactive knowledge base for the literature support of our network model (see links in data availability Section), while the main pathways that were included are listed below.

**Fig. 1:**
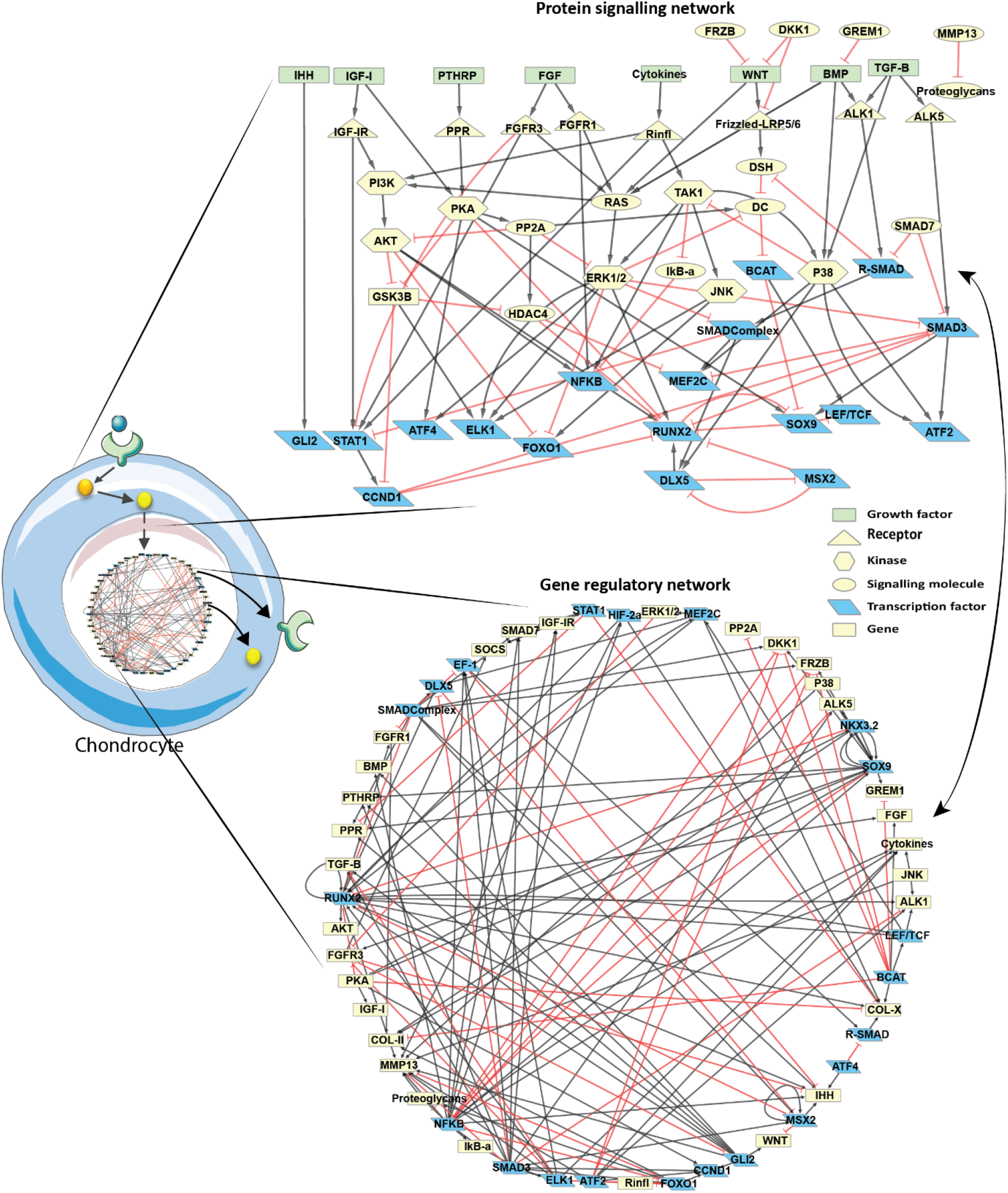
Influence map of the signaling and gene regulatory networks (GRN) implemented in the model. Red, T-ended arrows represent inhibitory influences and black arrows represent activating influences. On the signaling side, growth factors and pro-inflammatory cytokines are represented by green rectangular nodes, receptors are yellow triangles, kinase proteins are yellow hexagons and other signaling proteins are yellow ellipses. Transcription factors (TFs) are represented in blue both in the signaling network and the GRN. In addition, target genes are represented by yellow rectangles in the GRN and TFs might also be targets of other TFs in the GRN. Each biological component is represented by a gene in the GRN and a protein in the signaling network, except for nodes that are not involved in one of the layers (e.g. NKX3.2 only plays a role in the GRN while COL-II and COL-X do not have upstream or downstream influences in the protein signaling network)

Unless stated otherwise, all components of the network and of the *in silico* model are referred to with names in upper cases that designate the numerical variables, which represent neither the protein nor the gene but a product of both. Therefore, neither the gene nor the protein official nomenclatures are used. A list of correspondence between the numerical variable’s names and actual mouse gene names is available in **Table S2** and reported in the Cell Collective interactive networks, for information.

The regulatory pathways represented in the network include the canonical and non-canonical WNT BMP pathways, the PThrP and IHH pathways, as well as the IGF-I, FGFs and TGF-B pathways since they are all reported to play a role in chondrocyte fate decisions (*9*, *23*). The influence of pro-inflammatory cytokines, such as interleukine 1Beta (IL-1B) or Tumor Necrosis Factor Alpha (TNF-A), was summarized through a node labelled ‘cytokines’, in the network (**Fig. 1**). That node is able to signal through a single receptor (labelled ‘Rinfl’) that can activate well known downstream pathways such as the phosphatidylinositol 3-kinase (PI3K)/AKT axis, the nuclear factor kappa B (NFKB) pathway and mitogen-activated protein kinases (MAPK) pathways. These MAPKs in question include extracellular signal-regulated kinase 1&2 (ERK1/2), c-Jun N-terminal kinase (JNK) and P38. For each of the introduced pathways we represented the downstream signaling cascades and known transcription factors as well as their target genes in the nucleus. Examples of important transcription factors that were included are the IHH signal tranducer GLI2, the signal transducer and activator of transcription (STAT1), the transcription factor 7 (TCF), the myocyte enhancer factor 2C (MEF2C) as well as SRY-Box Transcription Factor (SOX9), a marker of differentiated healthy chondrocytes, and runt related transcription factor 2 (RUNX2), a hypertophy marker. All pathways in the model are strongly interconnected as shown by **Fig. 1.**

In total, there are 60 biological components in the network, which are listed in **Table S2**. Each component accounts for a different biochemical entity including 8 growth factors, 8 receptors, 20 transcription factors, 4 ECM proteins and 20 signaling molecules of another type. Combining the signal transduction and the gene regulatory networks into one connected network, leads to a total of 264 direct or indirect biochemical interactions. Each node has on average 7.2 direct neighbors (i.e. directly connected node) and the average shortest path (i.e. smaller number of edges) to connect two nodes is 3.01. Hence, all nodes may virtually influence others by means of a couple of intermediary components, supporting the need to represent and study that network mathematically to conclude on key controllers.

### Learning from transcriptomic OA data by complementing the mechanistic network with data-inferred interactions

We complemented the knowledge based GRN with data-driven network inference, allowing for identification of *de novo* regulatory links, introduction of previously unstudied or undiscovered interactions and the reduction of the bias related to human literature curation. To that end, we generated an informative cross-platform merged dataset on mouse osteoarthritic cartilage. It was composed of 109 samples coming from 6 microarray experiments from which we selected a subset corresponding to the expression profile of 41 genes (see Method and full list in **Data S1**). The selected genes were the ones present in or closely related to the biological factors from the mechanistic model. The purpose was to identify potential regulatory interactions among those genes of interest only (see Method). The sub-dataset was normalized and corrected for batch effects originating from the differences in microarray platform technologies, as described in the Method section. Our data indicate that the gene expression distribution was correctly normalized among the different microarrays (**Fig. 2A**). The principal component analysis (PCA) showed the inter-array variance was strongly reduced after merging and correction, thereby highlighting successful removal of the batch effect. As a quality control, we applied an unsupervised clustering method to the merged dataset to evaluate whether the biological information was still maintained after merging and correction. The resulting heatmap features the clustering of the 109 samples as well as their prior annotations as “OA” and “WT” (for wild-type or control samples) **(Fig. 2B)**. See Method and **Data S1** for annotations definition. While splitting the clustering result in three groups, we could identify an OA-like group and a WT-like group as well as a WT-like sample clustering alone. Based on these categories, 81% of the OA tagged samples were correctly identified in the OA group whereas 56% of the WT-tagged samples were correctly located in the WT group. These accuracy levels are similar to the ones achieved when clustering each of the individual microarray experiments separately, before batch effect removal (**Fig. S1**). We concluded that, despite the diversity of the technical platforms used in the assembled dataset, most of the variance was not due the arrays’ underlying technology but rather due to the biologically meaningful information usable as input for GRN reconstruction.

**Fig. 2:**
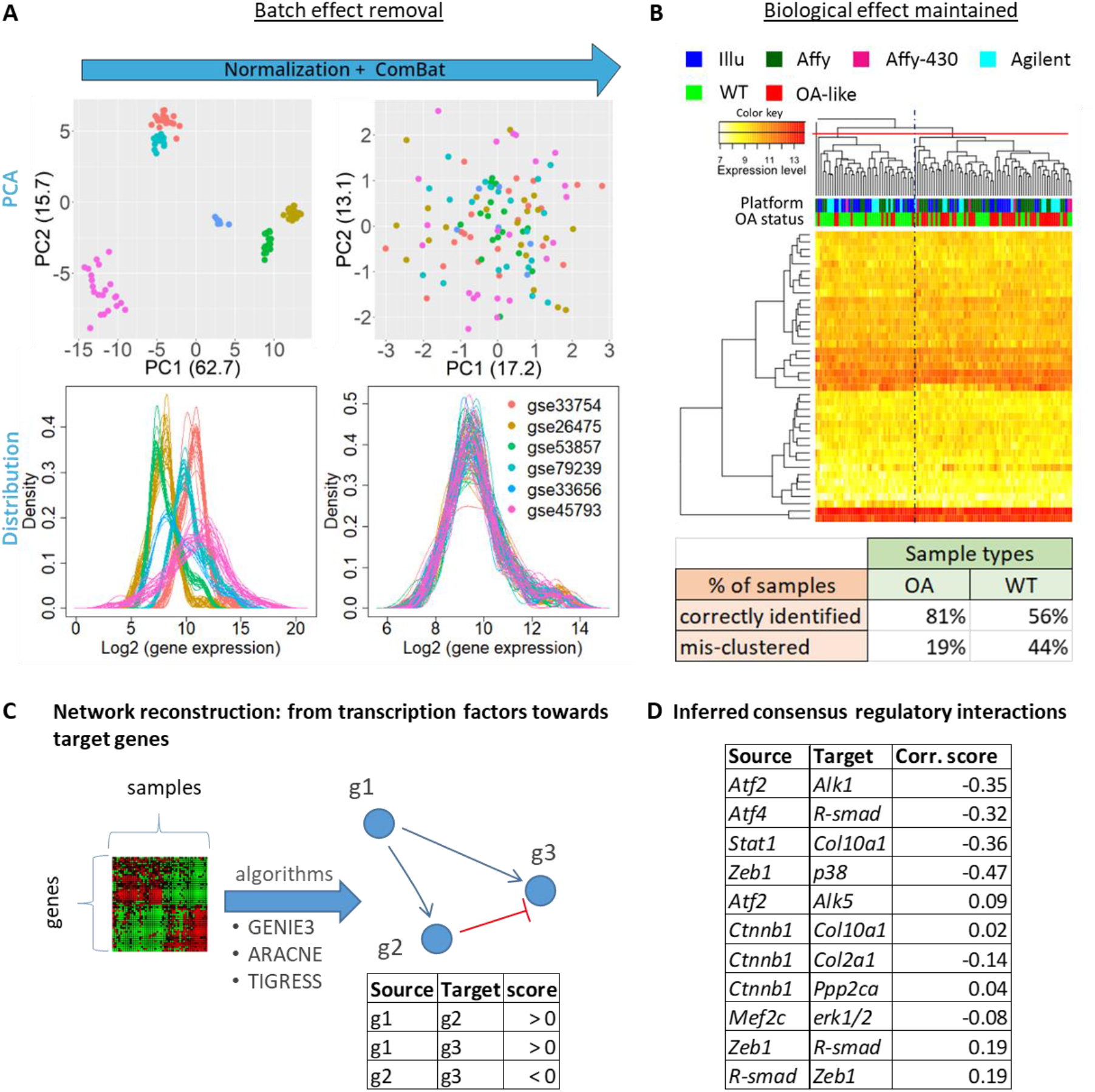
Micro array integration for GRN inference. **(A)** Assembling and correcting the microarray sub-datasets. PCA plots and gene expression distributions of the assembled dataset colored by arrays (GSE-number), before and after quantile normalization and batch effect correction with the Combat algorithm. **(B)** Unsupervised clustering with the Euclidian complete method highlights that the samples do not cluster according to the technological platform but rather according to the OA status of the samples. When splitting the hierarchical tree into three branches, a mostly OA group and a mostly WT group stand out. The table summarizes the percentage of true OA (resp. WT) samples correctly grouped in the ‘OA-group’ (resp. ‘WT-group’) highlighting a clustering accuracy in line with what is obtained for the individual sets before correction. **(C)** Starting from a matrix of gene expression as input, algorithms compute possible regulatory interactions and output a list of possible transcriptional regulatory interactions from a gene coding for a transcription factor to another gene. ‘g1’, ‘g2’ and ‘g3’ denote gene1, 2 and 3, respectively. **(D)** List of interactions inferred with the merged OA dataset and integrated into the mechanistic model. Inference was run with three algorithms, solely interactions that were present in the results of the three algorithms were reported and integrated. An interaction was considered present for one algorithm if its score was higher than a threshold defined as the difference between the mean and standard deviation of all scores. Corr.score is the spearman correlation coefficient, computed solely to define the interaction sign (activation if positive, inhibition if negative).

In order to complement the GRN with new data-derived hypothetical interactions, we inferred genetic interactions out of this newly assembled dataset. Directed edges between nodes were added in the GRN to account for newly inferred regulatory interactions, as represented in **Fig. 2C**. We inferred regulatory interactions directed from TFs towards target genes by using three different algorithms to avoid algorithm-specific bias on the inferred edges (see Method) (*24*). Only interactions predicted by all the three algorithms were implemented in the gene regulation layer of the mechanistic model. In addition, the sign of the Spearman correlation score allowed us to define whether the interactions were genetic activations or inhibitions (**Fig. 2C**). The inferred interactions are reported with the correlation factors in **Fig. 2D.** Note that the machine learning methods predicted some TFs as descriptor of a given target gene even for pairs that had not a high correlation score (**Fig. 2D**). This showed that the ensemble of algorithms we used went beyond the simple correlation for inferring genetic interactions, relying on the concept that some form of covariation is implied by a causal relationship. The inferred interactions were included in the GRN part of the model for all subsequent analyses presented in this study.

### The computational model successfully recapitulated two chondrocyte phenotypes and physiologically relevant behaviors

We translated the aforementioned regulatory network into mathematical equations in order to develop an executable numerical model of the articular cartilage chondrocyte. We used a semi-quantitative additive modeling formalism with priority classes as it allows to study large networks without requiring much information on kinetics parameters. Each node takes on a continuous value in the interval [0,1], representing the global functional activity of that node, defined as the multiplication of the gene expression level and the protein activation potential. That way, a protein cannot exert its function on downstream targets unless it is both expressed at the genetic level and activated/not blocked at the post-transcriptional level (global functional activity > 0) (see Method section, **supplementary method SM1** and **Table S1**).

With the above described numerical chondrocyte model, we studied the system free of fixed external cues to identify all potential mathematical stable states that may emerge naturally. These stable states are also called attractors or genetic modules and they equate potentially existing cell phenotypes (see definition of attractors in **Table S1**). They were evaluated using methodologies similar to those used with logical models (*25*). Any initial state inputted into the system of equations is like a set of external stimuli that would trigger signal transduction inside a cell, eventually leading to a specific cell state. By randomly initializing the *in silico* model (see method for Monte Carlo analysis) we were able to explore possible model outcomes and we observed three emerging attractors. Each of these attractors had a unique global activity profile for the 60 components as reported in **Fig. 3A**; the details of the protein activation and gene expression level for each component is available in **Data S2**. Two of the attractor states that we found were biologically relevant, meaning they were comparable to existing chondrocyte phenotypes identified based on known cell state biomarkers such as type II and type X collagen (COL-II and COL-X, respectively) , RUNX2 or SOX9. We identified one of the attractors as a normal healthy articular chondrocyte since markers such as SOX9, NKX.3.2 and COL-II were strongly active and expressed while RUNX2, COL-X and matrix metalloproteinase 13 (MMP13) were inactive or not expressed. In addition, the inflammation-associated nodes had low activity. That correlates with what is known of real chondrocytes during homeostasis. The second attractor corresponded to a hypertrophic-like chondrocyte since RUNX2, COL-X and MMP13 were present or active (i.e. global activities equaling to 1) while SOX9 was not and COL-II and the proteoglycans were degraded and/or not expressed (functional activities nearly zero). In addition, the WNT and inflammation related pathways were active (e.g. WNT =1, DC=0, βcatenin = 1, Cytokines =1, NFκB = 1, TAK1= 0.76). The third state we found had nearly all protein activities close to zero, consequently, we couldn’t associate it with any specific phenotype as it was more likely a trivial mathematical solution. We named it the ‘None’ attractor as it was neither healthy nor hypertrophic **Fig. 3A**.

**Fig. 3:**
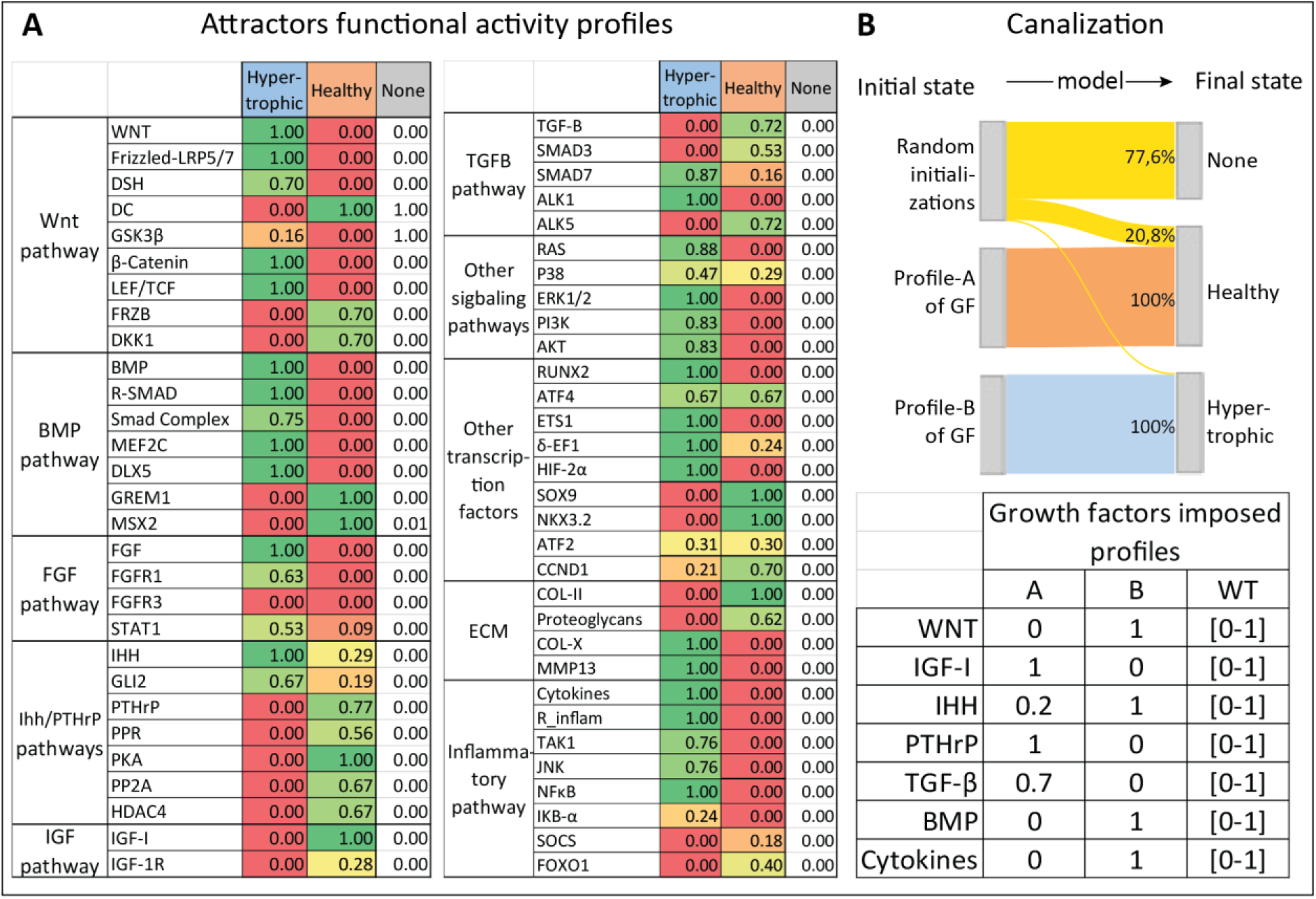
Predicted chondrocyte profiles and canalization for the three emerging states during Monte Carlo analyses. The Monte Carlo analyses consist in sampling 10.000 random initial states for the variables, with the possibility to impose constraints (similar to external biological cues). **(A)** The Monte Carlo analysis without constraints highlights the existence of three final states (i.e. attractors) with different activity profiles (in columns). The global activity of a protein is presented in this table as the product of the predicted gene expression and the protein activation level. The table only reports the global activity; a complete table including the gene expression and protein activation levels is available in **Data file S2**. Rows represent the variables of the model and are grouped by pathways or functional groups. **(B)** Specific activity profiles were imposed to the growth factors, as reported in the table, while variables other than growth factors were initialized randomly. Profile A represents a possible healthy environment. Profile B represents a more pathological environment. The ‘Random’ column indicates the case without constraints in which initial activities were randomly sampled within the interval [0,1]. In the Sankey diagram initial states are on the left and final destination states (i.e. attractors) on the right. Strips indicate the percentage of initializations (among the 10.000) that reached each of the possible attractorsnduring the Monte Carlo without constraints (‘Random initializations’).That percentage is also reported for the Monte Carlo with constraints (profile A & B).

The number of random initializations reaching an attractor during the Monte Carlo simulation gives a sense of the overall attractor’s size (attractors are reported in **Fig. 3B)**. Most of the random initial states led to the ‘None’ final state, reflecting the fact that most of these random initial values might very well be nonsensical from a chondrocyte biology perspective. In addition, about 21% of the initializations led to the healthy state and about 2% to the hypertrophic state (**Fig. 3B**). These attractors are the spontaneously emerging states arising from the proposed map of biochemical interactions.

We also sampled random initial states while fixing seven growth factors at values that were physiologically relevant in either a normal healthy or a disease environment (profile A and B of Fig.3B, respectively). Interestingly, when imposing the profile A to the growth factors during the Monte Carlo, the system only settled into the healthy state and when imposing profile B it settled into the hypertrophic state, thereby abolishing the ‘None’ state (**Fig. 3B**)

With the aim of assessing whether the model could successfully recapitulate relevant physiological behaviors, we established a tool enabling to test experimentally observable scenarios. As a first step to validate our model, we used this tool to test specific scenarios for which the expected outcome was known from literature or hypothesized from clinical observations. For instance, inflammation in the knee is one of the symptoms of OA and has been shown to be one of the drivers of cartilage degradation, possibly by pushing chondrocytes to undergo hypertrophy and produce matrix-degrading enzymes (*26*, *27*). For that reason, inflammation-related targets are subject to several investigations for potential OA therapies. Interestingly, *in silico* experiments with our model showed that blocking important transducers of inflammation such as the TGF-β-activating kinase (TAK1) or NFKB while activating the PTHrP related pathway could push a hypertrophic-like chondrocyte into transitioning towards a more healthy or anabolic state (**Data file S2**). Moreoever, other studies have reported that the TGF-β pathway had a protective effect against inflammation (*28*–*31*), a scenario we evaluated with our model too. *In silico* mimicking the presence of inflammation in a healthy chondrocyte by forcing several inflammation related pathways of the model to be set at their highest values led to 100% of transition towards the diseased hypertrophic state (**Fig. 4A**). This effect was partially rescued by concomitantly forcing the presence of TGF-β since 5.3% ±1.2 of the perturbations failed to exit the healthy state, thus, confirming *in silico* that TGF-β could have a protective effect against inflammation through the mechanisms present in the model. Nevertheless, the role of TGFβ in OA has been reported to be dual as this growth factor transduces signals in chondrocytes mainly via two receptors, the type I TGFβ receptor (ALK5) and the type II TGFβ receptor ALK1 (*32*). They are involved in different intra-signaling routes and depending on which receptor is activated, the downstream-activated signals would be rather anabolic (ALK5) or catabolic (ALK1) (*32*) and impact chondrocyte maturation differentially (*33*). Clinical observations reported that the ALK5/ALK1 balance decreased with age and in OA patients (*34*–*36*). *In silico* simulations with our model showed that, roughly, the rescue by TGFβ was lost when the ratio between the receptors was forced to be 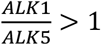 (**Fig. 4A** and **Fig. S2**). For higher values of ALK1, ALK1 activity could be as low as 0.86*ALK5 and still show the loss of the TGF-β protective effect (**Fig. S2**). This is in line with what was previously modeled (*28*), demonstrating that the decrease in that balance could explain why TGF-β loses its protective effect against inflammation in OA patients.

**Fig. 4:**
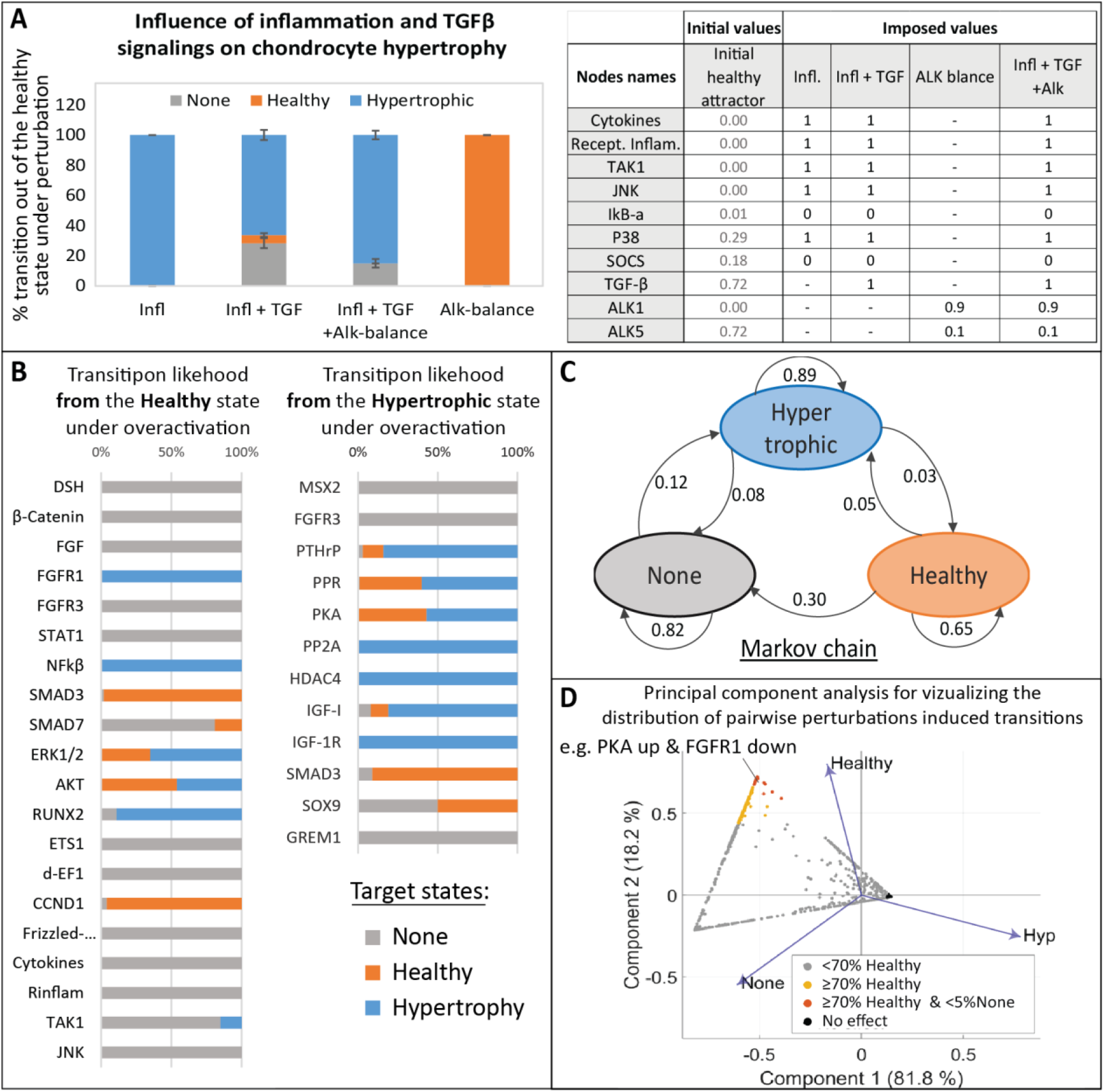
Study of the virtual chondrocyte state transition and *in silico* screening of target perturbations. **(A)** Relation between Inflammation and TGFβ and influence on the chondrocyte state. Perturbations are applied on the healthy attractor, bar height gives the average percentage of transition towardsone of the target states, error bar denotes standard deviation. ‘Infl.’ refers to imposed inflammation, ‘TGF’ refers to TGFβ over-activation and ‘Alk balance’ to the modification of the ratio between TGFβ receptors (ALK1 and ALK5). Conditions were mimicked as described in the table. ‘-‘ denotes no modification of the initial value. A transition from ‘healthy’ to ‘healthy’ means no transition. **(B)** All single node perturbations triggering a state transition from the Healthy (resp. Hypertrophic) state. **(C)** Markov chain providing the overall probability of transition from one state to another, under single node perturbations. Arrows indicate transitions from an initial state towardsa target state with the associated probability. Thus, the total probability of outgoing arrows for any state is 1.0. **(D)** PCA visualizing the results of the systematic screening of all possible combinatorial perturbations on a hypertrophic-like chondrocyte. Each dot represents one of the 7080 screened conditions. Principal components are computed based on the percentage of transitions towards the 3 attractors, reported as eigenvalues (blue arrows). Dot colors correspond to threshold in the percentage of transition towardsthe healthy state for potential OA therapies.

Together, these results show the ability of the articular chondrocyte *in silico* model to behave in a physiologically relevant way and predict emerging effects qualitatively, highlighting the pro- or anti-hypertrophic nature of biological components in specific conditions. The aforementioned tool was further implemented through an executable App (https://github.com/Rapha-L/Insilico_chondro) allowing users, such as biologists, to easily test hypotheses by performing *in silico* experiments on the virtual chondrocyte (see interface **Fig. S3**).

### *In silico* experiments on the modeled system predicted potential important nodes to control chondrocyte fate

We next decided to exploit further the model by studying the effect of all possible perturbations of each component. Starting either from the healthy or the hypertrophic-like state, variables were individually activated or inhibited in a systematic manner. Over-activation of the FGF receptor 1 (FGFR1) or NFKB were the most potent conditions to trigger a transition from the healthy towardsthe hypertrophic state. To a lesser extent, activation of the variables for ERK1/2 kinases, the AKT member of the PI3K/Akt axis, the RUNX2 transcription factor or JNK kinases also promoted this diseased transition (**Fig. 4B**). On the other hand, transition from the hypertrophic towards the healthy state was mainly triggered by forced activation of the TGFβ intracellular effector SMAD3 and partially brought about by activation of the SOX9 transcription factor, IGF-I and members of the PTHrP pathway (i.e. the protein kinase A (PKA), the PTHrP receptor (PPR) and PTHrP). This constitutes a state transition study that can be summarized in the Markov Chain representation (**Fig. 4C**) with the system’s overall probability of transitioning from one state to another under random single node perturbations. Interestingly, despite the small size of the hypertrophic attractor in the random canalization (**Fig. 3B**), the transition study shows that the total probability of transitioning out of that hypertrophic state under random single node perturbations is 0.11. So, in 89% of the cases, a single node perturbation will not affect this state. This means that even if the hypertrophic state is difficult to reach in the articular cartilage system, it is particularly robust to small single perturbations, and once the numerical system has reached that state, it is unlikely to escape from it, via transitioning to any other state, with a single targeting strategy. For this reason, we also systematically investigated the effect of all possible combinatorial perturbations of two constituents (or pairwise perturbation). The full screening amounted to 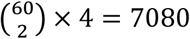 tested conditions per attractor. 94% of the variance was explained by the first two components in the PCA reporting the effect (percentage of transition to destination attractors) of the combinatorial treatments on a hypertrophic-like chondrocyte **(Fig. 4D)**. We searched for the most potent conditions to retrieve the healthy state from a hypertrophic chondrocyte (by requiring more than 70% of perturbations input transitioning to the healthy state and less than 5% to the ‘None’ state), which could point towards potential drug therapies for OA. Based on those results, the most efficient way to transition from a hypertrophic towards a healthy chondrocyte in the *in silico* model was with the up- regulation of SMAD3 in combination with activation or inhibition of numerous other targets such as inhibition of the inflammatory mediator NFKB, inhibition of the DLX5 transcription factor or activation of SOCS, a blocker of pro-inflammatory signals transduction (see complete list in **Data S3**). Activation of PKA/PPR in combination with inhibition of various targets, such as WNT or FGFR1, also seemed to decrease hypertrophy successfully in the model with 100% of transitions towards the healthy state (**Data S3**). Therefore, the PKA/PPR axis and SMAD3 seemed to be ‘enablers’ that could ‘unblock’ the *in silico* system, facilitating the effect of other relevant targeting treatments. Moreover, some predictions among the ones triggering more than 70% of transition towards the healthy state, in **Fig. 4D,** did not include the two aforementioned enablers. For example, the up-regulation of ALK5 in combination with the down-regulating ALK1, the two receptors of TGF-B in the model, gave more than 90% of transition towards the healthy state. Additionally, inhibition of the WNT pathway while activating ALK5 as well as activation of IGF-1 while activating the destruction complex (DC) involved in the WNT pathway allowed between 70 and 90% of transition towards the healthy state, to mention but a few (**Data S3**). So, the *in silico* model and associated screening algorithms enabled the prediction of pairwise targeting conditions with a good potential against chondrocyte hypertrophy.

### Newly predicted (combinatorial) treatments were validated *in vitro* for their potential to prevent hypertrophy

We used ATDC5s, a chondrogenic murine cell line able to undergo hypertrophy, in order to *in vitro* validate the *in silico* findings obtained from the model. We measured the activity of alkaline phosphatase (ALP), an enzyme typically secreted during hypertrophy and participating to ECM mineralization. It showed that there was a positive linear correlation between the level of ALP activity in the medium and the expression level of the hypertrophic gene *Col10a1* during the ATDC5 differentiation (**Fig. 5B**). Hypertrophy was further increased when supplementing the differentiation medium with Ihh, as expected (**Fig. 5B**). Indeed this growth factor is known for its pro-hypertrophic effect on ATDC5 (*37*). These results allowed us to evaluate ALP activity in the medium as a proxy for the level of hypertrophy, significantly increasing the throughput of the experimental set-up for testing small molecule treatments.

**Fig. 5:**
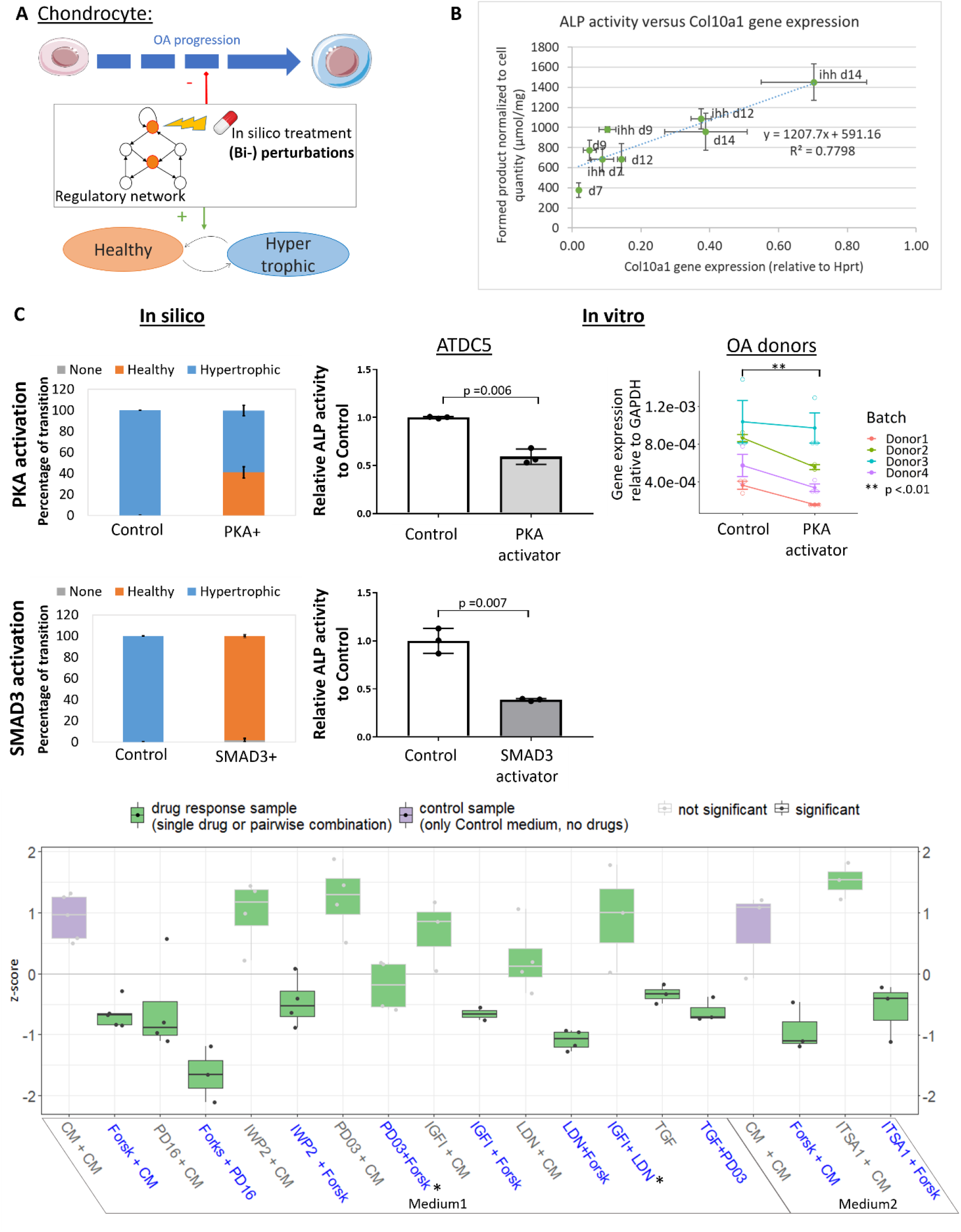
*In vitro* validation of *in silico* predictions on chondrocyte phenotype changes. **(A)** Concept of *in silico* identification of potential drug targets. **(B)** Secreted ALP activity, relative to DNA quantity, is positively linearly correlated to *Col10a1* gene expression during hypertrophic differentiation with and without Ihh treatment. Each point is the average of 3 replicates and bars denote standard deviation. **(C)** Effect of PKA or SMAD3 activation as measured *in silico* and *in vitro* in ATDC5 (N = 3 replicates, histograms show average fold change in ALP activity relative to control and bars are standard deviations, p-values are computed with one-tailed t-test and welch’s correction) and chondrocytes from OA donors (N= 4 donors with 3 replicates each, p-value is computed with one-tailed linear mixed effect model). *In silico* activation was performed by setting the nodes to their maximum value (1.0), *in vitro* PKA (resp. SMAD3) activation was performed with Forskolin 1μM (resp. Activin 100ng/ml) for 24h. **(D)** Single and combinatorial drug screening in ATDC5 with selected conditions based on *in silico* predictions. Boxplot of the series of conditions across independent replicates (z-scores of ALP activity fold change) with control conditions in purple. Conditions significantly lower than the control (combined p-value < 0.05) have dark grey borders and dots (Wilcoxon rank-sum test with BH correction and combined probabilities over independent runs). For each condition, dots are the average of biological triplicates, summary statistics are represented by a horizontal line for the median of independent experimental repetitions and a box for the interquartile range. The whiskers extend to the most extreme data point that is not <1.5 times the length of the box away from the box. Blue labels indicate potent conditions predicted by the *in silico* model, grey labeled conditions are added to the experimental set-up for information. CM stands for ‘control medium’, medium1 has 0.02% of DMSO and medium 2 0.035%.*indicates *in silico* predicted conditions that did not show a significant decrease of ALP activity *in vitro*.

This semi-high throughput system was used for validating the *in silico* predictions. *In vitro* experiments evidenced that treatments with Forskolin, an activator of PKA activity or with an activator of the BMP/SMAD3 pathway, were sufficient to prevent the increase in the activity of secreted ALP and thereby were sufficient to block hypertrophy in ATDC5s (**Fig. 5C)**. Additionally, Forskolin treatment decreased *COL10A1* gene expression when applied to primary human OA chondrocytes (p < 0.01) (**Fig. 5C**). Together these results corroborated the *in silico* predictions that activation of PKA was sufficient to block hypertrophic differentiation in chondrocytes (**Fig. 5C & D)**.

Additionally, several combinatorial treatments predicted by the *in silico* screening were tested in ATDC5 and compared to the corresponding individual treatments, to assess their efficacy. In **Fig. 5D**, ALP activities normalized to total DNA content are reported by means of z-score for all pairwise and single conditions. The ALP activity in the medium during hypertrophic differentiation was lower than the corresponding controls in 6 out of the 8 predicted combinations. The two conditions for which the strongest effect was measured were the inhibition of FGFR1 combined with the activation of PKA (‘Forskolin + PD161570’, p = 0.032) and the inhibition of BMP combined with PKA activation (‘LDN-193189 + Forsk’, p = 0.014). These two combinations seemed to generate an added effect compared to the single drugs. Contrary to *in silico* model predictions, treatment with exogenous IGF-1 combined with BMP inhibition (‘IGF-I + LDN-193189’, p= 0.105) or inhibition of ERK1/2 combined with PKA activation (‘PD0325901 + Forsk’, p = 0.264) did not show a significantly lower hypertrophic level, based on ALP activity (**Fig. 5D**).

Elaborating one of the conditions that showed the strongest response, PKA activation combined with FGFR1 inhibition, we compared the combinatorial effect with the corresponding single drug treatments. The combinatorial effect was greater than the one for either of the single drug, for both tested concentrations ratios (**Fig. 6A**). This suggests that activating PKA (resp. inhibiting FGFR1) would potentiate or enable the effect of FGFR1 inhibition (resp. PKA activation) by blocking or unblocking key pathways and maintaining the necessary constraints on the system. Dose curve relationships need to be established to confirm that hypothesis. Screening various values of functional activities of PKA and FGFR1 with the virtual chondrocyte and looking at the percentage of transition out of the hypertrophic state, showed that a minimal level of PKA activity (namely about 0.4 on a scale from 0 to1) was required to achieve any positive effect with this combinatorial treatment (**Fig. 6B)**. In addition, a gradient effect was observed suggesting that the lower the PKA activity, the more we needed to block FGFR1 to achieve an equivalent positive effect (**Fig. 6B**). *In vitro* validation confirmed this dose effect since the overall gradient shape was very comparable between the *in silico* and *in vitro* situation **(Fig. 6B)**. Comparing the diagonal (combinations of two drugs) to the single drug ranges also highlighted a likely synergistic effect in decreasing hypertrophy between the two drugs, at the tested 1:5 ratio. Taken together, these results support that targeting the regulatory mechanisms at multiple points might be necessary to maintain a physiologically healthy state for a chondrocyte experiencing hypertrophy inducing cues.

**Fig. 6:**
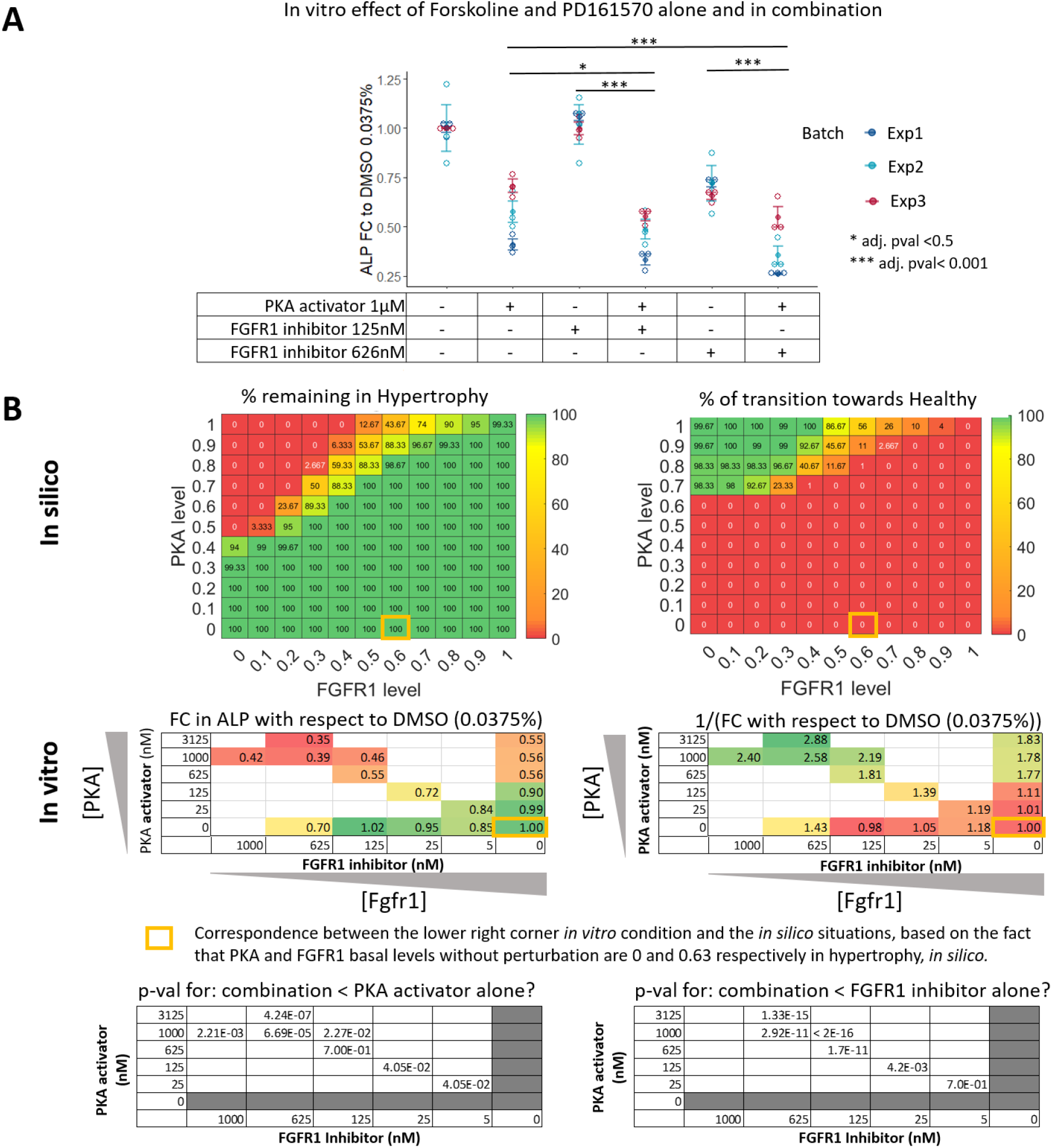
*In silico* v.s. *In vitro* dose response effect of PKA activation with FGFR1 inhibition. The most potent condition from the screening is investigated further for a potential dose effect. **(A)** Fold change (FC) in ALP activity, with respect to control, due to PKA activator (Forskoline, 1μM) or FGFR1 inhibitor (PD161560, 125nM and 625 nM) or the combination of both. **(B)** A range of values for PKA and FGFR1 imposed activities is screened *in silico* with 0 meaning no activity and 1 being the max possible activity. The percentage of transition remaining in the hypertrophic state or transitioning towards the healthy state are reported in the upper panels, the rest of the transitions goes to the ‘None’ state. In the middle panels, fold change (resp. inverse of fold change) in DNA-normalized ALP activity with respect to control DMSO in ATDC5 is reported for a range of Forskolin and PD161570 concentrations. The *in vitro* situation without drugs (yellow rectangle) would correspond to the basal level of PKA and FGFR1 in *in silico* hypertrophy but there is no one to one correspondence between the *in silico* and *in vitro* ranges. All *in vitro* results represent n=9 (3 bio-replicates in 3 independent experiments), p-values are computed on log-transformed data with a linear mixed-effect model, user defined contrasts (only combination versus corresponding single doses were compared), one-sided test and adjustment for multiple comparisons with the Holm’s method. The combinatorial treatment effects were greater than the ones for either of the single treatment both *in silico* and *in vitro*, for all concentrations in the gradient of dose relationships investigated.

## 3. Discussion

We report the construction of a mechanistic model of chondrocyte phenotype control in articular cartilage by combining a knowledge-based approach with a data-driven approach. We have leveraged decades of knowledge and data about chondrocyte regulatory pathways and osteoarthritis by integrating that information in a numerical predictive model. This model enables to recapitulate physiologically relevant observations and predict conditional effects resulting from intricate intracellular signalings. It offers the possibility to screen a large amount of (combinatorial) treatments and prioritize subsequent *in vitro* experiments for the identification of molecular drivers and drug targets in OA. The *in silico* targets perturbation screening and the experimental validation of our findings show the potential of *in silico* experiments to identify relevant *in vitro* experiments and refine the early drug discovery pipeline for OA. In particular, our study points towards a likely synergistic effect of PKA and FGFR1 targeting strategies to regulate chondrocyte hypotrophy. Several of our insights have implications both for the network modeling community and for cell and cartilage biology.

We have first built an interactive intracellular network as an online knowledge base and as a reference support for our numerical model. Most network-based models rely on prior mechanistic knowledge. Even current state of the art computational tools meant to reconstruct numerical models automatically from (high-throughput) data often require or offer the possibility to introduce prior knowledge (*38*). Therefore, there is high need for integrating and curating originally isolated pieces of knowledge, in a comprehensive way. However, biochemical information specifically related to cartilage and osteochondral systems are scarce in public and private pathway databases, in which cancer related cell types tend to be over-represented. The network we provide along with this study, details the prior mechanistic knowledge we have put in the model and that was predominantly chondrocyte or osteochondral cell type specific, with a focus on articular chondrocytes and osteoarthritis. In our opinion, it is valuable not only for cartilage and OA researchers but also for modelers as it can serve as a basis to derive other models to answer alternative questions.

Combining knowledge and data in a comprehensive network resulted in a systemic view of chondrocyte intracellular regulation. Many independent pieces of information have accumulated over the past decades and many databases have made curated pathways available. Most of the time in literature, these pathway descriptions stop after the (in)activation of the downstream transcription factors while the identity of the target genes downstream these pathways is left obscure. In this study, our strategy was to complement the knowledge based network graph with automatically inferred transcriptional regulations from transcriptomic data through the use of machine learning methods. This usually requires a large multi-perturbed dataset but in absence of such dataset for chondrocytes, we reconstructed one by merging various arrays. Our strategy is supported by a previous study, which reported that equivalent informative data could be successfully achieved by assembling naturally occurring and experimentally generated phenotypic variations of a given cell type (*39*). Even though relatively few inferred gene regulatory interactions were integrated in the mechanictis model due to the stringency of our selection strategy, we limited the risk of integrating false positive predictions. This data-driven approach allowed us to take advantage of automatic network reconstruction technologies, which are becoming more and more the standard in the state of the art (*40*, *41*).

The translation of that network into mathematical rules was then further studied computationally and enabled the prediction of the overall effect of each network component on the virtual chondrocyte. Our perturbation screening approach helped to discover influencer nodes based on the mathematical dynamics, similarly to what had been previously proposed for other diseases (*17*). Important to mention is that no information was put in the model that directly made one target prevail over the others in being pro- or anti-hypertrophic. The information that fed the mathematical model was about activating or inhibitory influences of one molecule on another molecule, a complex of molecules or a pathway although some edges in the network represented indirect links for the sake of simplification. The advantage of the discrete semi-quantitative mathematical formalisms we employed lies in its ability to reproduce qualitative dynamics using only the activating or inhibitory nature of interactions and additive rules, without any prerequisite about kinetics. This characteristic makes that method perfectly suitable for a large network such as the one of this study. Many biostatisctics and machine learning methods that make drug efficacy related predictions solely based on omics data still lack predictability and interpretability (*42*). In contrast, the numerical approach presented in this study has the advantage of providing mechanistic evidence supporting the predicted effects, thereby increasing the mechanistic interpretability.

Generally, the validation of an *in silico* model to answer questions in a specific context may be achieved in two ways: either by showing its ability to predict non-linear effects that were already reported in literature or by confirming predictions with new experiments. The relevance of our model to answer osteoarthritis-related questions was first confirmed by its ability to recapitulate earlier described behavior such as the changing role of TGFβ signaling in presence of inflammatory stimuli. A recent computational study reporting a quantitative (ODE-based) time dependent model of cartilage breakdown (*28*) provided more detailed and complexed kinetics underlying the dual role of TGF-β. The fact that our model could lead to equivalent conclusions shows that the more simplifying semi-quantitative formalism that we employed could suffice to capture non-straightforward effects, thereby supporting its credibility for subsequent predictions.

The Monte Carlo analysis of the system without constraints showed that the healthy state was more easily reached than the hypertrophic one. This corroborates with the situation for normal articular cartilage in which chondrocyte hypertrophy does not spontaneously occur unless the homeostasis is disturbed (*43*). A great part of the state space was occupied by the ‘None’ attractor that is most likely a trivial solution towards which the system converges when an initial state or a trajectory is too far away from a feasible biological state to meet all the constraints imposed by the equations. In such a situation the trivial ‘zero’ solution is more easily reached. An analysis of the ensemble of initial states reaching the ‘None’ state, could possibly spotlight initial states that are unlikely to happen in an *in vivo* physiological environment. Conversely, the analysis of canalization under imposed growth factor profiles have suggested that restricting ourselves to initializating growth factor’s values within biologically ‘feasible’ ranges can decrease prevalence of the ‘None’ state, although the boundaries of those feasible ranges could be further explored. Nevertheless, in the scope of finding potential therapeutic targets, we are less interested in the random canalization of the states than in their robustness to perturbations and capacity of transitioning.

An important result of this study is the highlights that chondrocyte phenotypes are not much sensitive to small, single factor perturbations. Even though, the resistance of the current *in silico* model to single factor perturbation might, in part, be due the omission of some parts of the real world system’s complexity, this trait of robustness to small environmental variations is often considered as a fundamental and ubiqituous trait of biological systems (*44*). This traits allows systems to function in noisy environements (*45*) and sytems biologists have theorized that disease may establish its own robustness, in some cases (*46*). In line with that, the Markov chain in our computational model indicates that, for articular chondrocytes, the probability to change phenotypes once it has been reached is rather low. Indeed, due to the very intricate interplay of molecules, it is likely that some pathways play redundant roles and that several factors should be targeted simultaneously to ‘unlock’ the system. Moreover, our *in silico* pairwise perturbation screening confirms that targeting at least two proteins at the same time increase the chance to unlock the hypertrophic commitment. Overall, as its *in vivo* counterpart, the *in silico* articular chondrocyte is unlikely to display hypertrophic signs in normal conditions or even sometimes under inducing treatment (*10*), but once the hypertrophic transition has been initiated, it is rather difficult to escape that fate.

Another important outcome is the experimental validation of the model predictions on previously unreported conditions. In practice, as reverting a hypertrophic chondrocyte back to a healthy state has never been observed experimentally, we hypothesize that the *in silico* predicted conditions are, at least, more likely to block hypertrophy. The *in vitro* results that we present here support this hypothesis and open new routes for further testing the suggested combinatorial conditions. Especially, the combination of PKA activation with FGFR1 inhibition is highlighted as good candidate treatment by our integrated *in silico*-*in vitro* approach. Activation of the PThrP pathway, to which PKA belongs, has already been reported to be rather anti-hypertrophic for growth plate chondrocytes (*47*). Similarly, genetic inhibition of the Fgfr1 gene in mouse knee cartilage has been shown to attenuate the degeneration of articular cartilage in mice (*48*). However, to our knowledge, this combination has never before been investigated nor reported for its synergistic potential against hypertrophy, cartilage degradation or OA.

Validating *in silico* predictions for drug target discovery experimentally is a challenging task. This is especially true when numerical high-throughput screenings, enabled by high computing power, generate a large amount of predictions. We leveraged the evaluation of secreted ALP activity in ATDC5 cell line as a semi-high throughput read-out for validating the *in silico* predictions. Indeed, it served as a convenient experimental system to assess hypertrophy modulation under many screened perturbations (*49*). Even if that cell line is of mouse origin and cultured in isolation from a physiological environment, we propose to use it within an *in silico*-*in vitro* pipeline that would act like a funnel for prioritization of candidate drug targets. As shown in our study the main hits can then be further evaluated in more detail using human OA chondrocytes.

We acknowledge that our study has some limitations. A first drawback is the absence of a gold standard to assess the precision of the data-driven network inference. To mitigate this, we used a consensus approach by only integrating predictions made by three different machine learning methods. This reduced the amount of interactions we would integrate but it alleviated each method’s weakness and reinforced the strengths of the predictions, as previously proved (*24*). Secondly, the type of mathematical model we employed comprises almost no numerical parameters but the main one, the saturation constant, was assigned an arbitrary value based on a previous study (*20*). This constant determines how fast a protein activity or gene expression can saturate to the maximal value depending on the amount of excess positive and negative upstream interactions. It intervenes in the weight of interactions and changing its value might slightly affect the influence of the network’s constituents on the system. Finally, we did not experimentally verify the target specificity and dosing regimen for the small molecules employed for our *in vitro* validation since it was out of the scope of this study. However, those small molecules were selected based on their previously reported and well-known *in vitro* action on our targets of interest.

This study is a proof-of-concept that an *in silico - in vitro* integrated approach can suggest single and combinatorial target perturbations affecting the hypertrophic transition and help to prioritize therapeutic targets for OA therapy development. In that sense, our model offers the possibility to draw conclusions on the pro- or anti-hypertrophic nature of biochemical pathways and targets based on strict mathematical rules describing the intricate network connectivity. We are convinced that this type of approach can guide the process of therapy development from basic understanding to target selection early on in the drug discovery pipeline while reducing time and cost of experiments as well as the use of animal models in early stages of drug discovery. Furthermore, investigating the effect of a new target that was not present in the current model should be possible by solely informing the model on how the target interacts with and connects to the rest of the network. Typical information on the nature of the upstream activators and inhibitors of the protein’s functionality, the nature of downstream proteins modulated by the target under scrutiny as well as information about its transcriptional regulators, from DNA binding assays for instance or reverse engineered from data would be needed. Ideally, this information should be as exhaustive as possible based on current state of the art knowledge, while hypothetical connections could be investigated and compared based on the simulated target’s effect. Finally, as scientific research is making progress in the identification OA disease subgroups based on molecular markers and clinical phenotypes (*50*, *51*), we foresee data-informed mechanistic models can become more and more patient-type specific. For instance, such knowledge-based network model could serve as a prior and be further optimized with engineering approaches, adjusting the network topology and/or interaction weigths, in order to fit chondrocyte baseline profiles of typical patient subgroups. Then, the resulting network models and the effect of targets’ perturbations could be compared across the different patient type-specific models.

## 4. Material and Methods

### Network construction

The knowledge-derived networks were built by incorporating fine-grained mechanistic knowledge about signaling pathways and transcriptional regulations. A previously published model of chondrocyte differentiation in the growth plate (*21*) was used as a basis and was adapted and completed through literature curation of decades of knowledge about articular chondrocyte and osteoarthritis,. Reference sources for the experimental evidence (binding assays, clinical observations, etc..) were mostly from mouse and human origin. References used chondrocytes (or related cell lines) and predominantly involved direct protein or promoter binding information, curated pathways or observed phenomenon in cartilage during homeostasis and disease. All references and mechanism descriptions are available through the interactive networks in the online platform Cell Collective (http://www.cellcollective.org) (*22*). See the links for the chondrocyte knowledge bases in the ‘Data availability’ section. Additionally, some gene regulatory interactions were automatically identified with machine learning algorithms using gene expression multi-perturbed data as input. Such informative dataset was achieved by merging six published microarray datasets of mouse articular cartilage (GSE26475, GSE33754, GSE79239, GSE33656, GSE53857, GSE45793). Each dataset had a control or wild type group and an OA-like group, which was either using genetically modified mouse spontaneously developing OA or a DMM-induced OA mouse model. Importantly, the initial annotations provided by the authors were not necessarily ‘OA’ or ‘WT’ but rather experiment-specific annotations such as ‘DMM’ or ‘SHAM operated’. Therefore the binary annotation as OA-like or WT-like samples was established by hand for the purpose of this study according to the original data annotations published on GEO, as reported **Data S1**. The datasets were merged applying an in-house developed pipeline adapted from previously published methods (*52*). Briefly, the datasets were preprocessed first in a platform-specific way prior to assembly as a merged dataset. For the purpose of this study, a sub-dataset was created by restricting ourselves to the genetic profile of 74 genes of interest, being the ones present or closely related to the biological factors from the mechanistic model as listed in **Data S1**. 41 genes, out of the 5470 from the full dataset, matched our list of interest, based on the Ensembl IDs, and the associated genetic profiles constituted the final sub-dataset (41 genes (or variables) x 109 samples (or observations)). This dataset was quantile normalized and the batch effects were removed through the ComBat algorithm based on Bayes methods (*53*).

Inference was performed for regulatory interactions within the aforementioned subset of genes. This was achieved employing three different algorithms, being ARACNE, TIGRESS and GENIE3 (*54*–*56*); the final retained network was a consensus network of the three algorithms. Typically, an interaction between a transcription factor and a gene was kept if it was present in the three methods’ results. An interaction was called present for a method if the interaction score was higher than a certain threshold for that method. This threshold was set to *m* − *σ*, where *m* is the average score for the given method and *σ* is the standard deviation of the scores.

Unless specified, all the aforementioned microarray data analytics were accomplished using the R computational environment (v.3.2.2).Topological parameter analysis of the final network was carried out with the Network Analyzer plug-in of the Cytoscape software v3.7.2 (https://cytoscape.org/) (*57*).

### Implementation of the mathematical model and dynamic analysis

The information contained in the network was translated into mathematical equations through an additive formalism with 2 priority classes to distinguish between fast and slow reactions, the importance of which was repeatedly highlighted (*58*–*60*). This additive formalism resembles the Boolean threshold networks (*61*) and was previously implemented with priority classes as described by Kerkhofs and Geris (2015). In this formalism, the nodes or biological component were represented by variables for which the values evolve over pseudo-time steps. The model is semi-quantitative since the nodes or variables could take on a continuous activity values between 0 and 1. The evolution of variables (proteins or genes) was defined by the sum of the upstream activating variables and the subtraction of the upstream inhibitory variables from the network (see definitions in **Table S1**). Biological interactions could happen at two time scales, reflecting the priority classes: reactions related to slow biological processes such as gene expression, mRNA or protein production, were referred to as slow reactions (lower priority) and those related to fast processes such as protein activation or inhibition, were referred to as fast reactions (higher priority) (see **Table S1** for the definition of fast and slow reactions & variable). A formal description of the mathematical system underlying the model as well as the full list of equations are available in **supplementary method SM1**.

The asymptotic solutions were evaluated with a Monte Carlo simulation procedure, similar to methods employed for logical models (*25*, *60*). When running a simulation (also see section below on Monte Carlo analysis), an initial value in the interval [0,1] was assigned to each variable. Every simulation step, the sub-variables (see definition in **Table S1**) were updated asynchronously according to the rules given in the equations and following the priority classes, in such a way that fast reactions were always updated before the slow reactions. The order in which variables were updated within a priority class was random, hereby recapitulating the stochasticity inherent to any biological system. See **Fig. S4 and S5** for graphical explanations about the algorithm and simulation scheme on a reduced illustrative example network. A stable state (definition in **Table S1**) was reached whenever the next iteration step did not bring any change for any of the variables up to a tolerance 10^−2^. In other words, when initializing the system at a random point, it was considered converged when the relations detailed by the system of equations were fulfilled up to a tolerance of 10^−2^. Thanks to the stochasticity of the model, the same initial point could lead to different types of stable states. Therefore, all computational results of this paper were computed 3 times and standard deviations were evaluated. All implementations and simulations were carried with the MathWorks® suite, MATLAB (2018b).

### Monte Carlo analysis and estimation of attractors (or stable states)

The nature and size of the stable states, given the regulatory network provided in the equations, was estimated by a Monte Carlo procedure. In short, all variables were initialized 10.000 times with random values in the interval [0,1]. The amount of initializations reaching each final stable state were computed and reported in terms of percentage of initial states. This number gives a sense of the stable states’ size for the unperturbed system, i.e. without constraints (*25*, *60*). Then, we performed two other Monte Carlo canalizations in which all variables were randomly initialized, except for seven growth factors that were fixed at values being physiologically relevant in either a normal healthy or a disease environment (profile A and B of Fig.3B, respectively). The networkD3 package from R was used to produce the Sankey diagram for the visualization of the canalization results.

### *In silico* target perturbations

By essence, the attractors are stable, meaning that variables cannot evolve anymore. These states may however be escaped by forcing, computationally, the value of one or several variables to change. Such a perturbation was imposed for a fixed amount of computational iterations, after which the system was left to evolve freely, thereby accounting for the fact that chemical treatments affect biological systems for a finite period of time. The duration of the perturbation was set to 1000 time steps as the perturbed state did not take more than 200 time-steps to be reached, on average. Imposing a perturbation on a stable state forces the system to evolve again, following rules imposed by the equations, and eventually settle down in the same initial or a new attractor (see convergence description above).

The different *in silico* scenarios or treatment experiments that were tested amounted to perturbing one or several variables/nodes, from the healthy or the diseased hypertrophic state and assessing the effect of that perturbation on the state stability. Variables were perturbed by forcing their global activity value to be 0 or 1 for inhibition or activation respectively. Imposing intermediary values between 0 and 1 was also done for some specific questions in which extreme values would be unlikely, such as varying the ratio between different membrane receptors. Each perturbed conditions was imposed starting from the relevant initial stable states (healthy or diseased) and the nature of the final state to which the perturbation led after simulation was documented. We considered that the tested perturbation triggered a state transition when the final state was different from the initial one. Given the stochastic nature of the model, the same perturbation could trigger a different outcome if simulated a second time, therefore the same perturbation was repeated 100 times and we reported the percentage of transition towards each of the possible target states. Standard deviation in the percentage of transition was assessed by repeating that experiment 3 times.

Due to the computational cost associated with the systematic screening of all possible pairwise perturbations, (for each pair there are 4 possible pairwise conditions to be tested either from the healthy state or from the hypertrophic state. The independent simulations were run in parallel using high performance computing infrastructure of the KU Leuven (Vlaams Supercomputer Centrum). Afterwards, we automatically selected combinatorial conditions for which at least 70% of the perturbations triggered transitions from the hypertrophic state towards the healthy one. Among them, we focused on those conditions with more than 70% transition towards the healthy state but less than 5% towards the ‘None’ state as a first discriminatory factor. Conditions were further selected for their druggability and their ease to be tested in a simple *in vitro* system. For instance, conditions involving the modulation of transcription factors were not considered for the *in vitro* validation since, no small molecule treatment could directly affect transcription factor activity.

### Testing treatments in vitro with ATDC5 culture

The validation of our *in silico* predicted treatments required *in vitro* testing with small molecules. This was performed with ATDC5, a mouse chondroprogenitor cell line obtained from the Riken Biological Ressource Center. Cells were cultured in 2D in proliferation medium containing DMEM/F12 (ThermoFisher, UK), 5% Fetal Bovine Serum (Biowest, Belgium) and 1% antibiotic/antimycotic (Gibco, ThermoFisher Scientific). Chondrogenic differentiation was induced by plating the cells at 6,400 cell/cm² in proliferation medium for 24h, followed by changing the medium to differentiation medium, being proliferation medium supplemented with 10 μg/ml insulin (Sigma-Aldrich), 10 μg/ml transferrin (Sigma-Aldrich) and 30 nM sodium selenite (Sigma-Aldrich). Cells were incubated in a humid environment at 37°C, 5% CO2 and differentiation medium was refreshed every other day for the first 10 days, and every day after the 10^th^ day, for longer experiments (i.e. Fig. 5B). Supernatant medium was taken for ALP activity assay and cells were harvested for DNA quantification (0.05% Triton-X reagent). ALP activity was reported relatively to the total DNA quantity to alleviate potential variation in cell number.

To study the correlation between *Col10a1* expression and secreted ALP activity, cells were differentiated for 14 days with or without Ihh supplement (150ng/ml, R&D Systems Europe LTD). The cells were harvested for RNA isolation on day 0, 7, 9, 12 and 14 during differentiation (TRIzol reagent; Thermo Fisher Scientific), in addition to the ALP acitivity assay and DNA quantification.

To assess the effect of small molecules and growth factors treatments on hypertrophic differentiation, cells were treated on day 8 of ATDC5 chondrogenic differentiation for readout at day 9. Cells were treated with one or a combination of the following compounds: Forskolin (1μM, Axon Medchem), Recombinant Human/Mouse/Rat Activin A Protein (100ng/ml, R&D Systems), Recombinant Mouse IGF-I/IGF-1 Protein (10ng/ml, R&D Systems), Transforming Growth Factor (TGF)β1 (10ng/ml, PreproTech), PD0325901 (1μM, Axon Medchem), PD161570 (1μM, Axon Medchem), ITSA1 (50μM, Chembridge), LDN-193189 (0.5μM, Axon Medchem), LY294002 (20μM, Axon Medchem) and IWP2 (2μM, Stem cell technology). ALP activity in treated conditions is expressed in terms of fold change with respect to the control medium with appropriate amount of DMSO, which was used as a solvent for most small molecules. Four types of control media were used throughout this study due to sparse solubilities of the compounds. The control medium was with 0.02% DMSO (Medium1) for most treatments, with 0.1% DMSO (Medium2) for the ITSA1 related conditions, without DMSO) for the ActivinA treatment (**Fig. 5),** and with 0.0375% DMSO for the Forskoline/PD161570 synergy study and dose screening (**Fig. 6**).

### Validating treatments effects in primary human chondrocytes culture in alginate beads

Human articular cartilage was obtained with implicit consent as waste material from patients undergoing total knee replacement surgery. This protocol was approved by the medical ethical committee of the Erasmus MC, University Medical Center, Rotterdam, protocol number MEC-2004-322. To isolate chondrocytes, cartilage chips were subjected to protease (2 mg/ml, Sigma Aldrich) for 2 hours followed by overnight digestion with 1.5 mg/ml collagenase B (Roche Diagnostics, Switzerland) in Dulbecco’s Modified Eagle’s Medium (DMEM) high glucose supplemented with 10% fetal bovine serum. Single cell suspension was obtained by filtrating the cellular solution by a 100 μm filter. The isolated chondrocytes were expanded in monolayer at a seeding density of 7,500 cells/cm2 in DMEM high glucose supplemented with 10% fetal bovine serum, 50 μg/ml gentamicin and 1.5 μg/ml fungizone (Gibco, Grand Island, NY, USA). Upon approximately 80% confluency cells were trypsinised and reseeded at 7,500 cells/cm2. Cells were used for experiments after three passages. Redifferentiation of articular chondrocytes was performed using a 3D alginate bead culture model. For preparation of alginate beads, chondrocytes after three passages in monolayer were re-suspended in 1.2% (w/v) low viscosity alginate (Kelton LV alginate, Kelko Co, San Diego, CA, USA) in 0.9% NaCl (Sigma Aldrich) at a concentration of 4 × 106 cells/mL. Beads were made by dripping the cell-alginate suspension in 105 mM CaCl2 (Sigma Aldrich) through a 22-gauge needle. Beads were washed with 0.9% NaCl and DMEM low glucose. Beads with a size that deviated from the average after a visual inspection were not included in the experiment. Re-differentiation of chondrocytes was performed in a 24-well plate (BD Falcon) for two weeks in 100 μL/bead DMEM low glucose supplemented with 1% ITS fetal (Biosciences), 10 ng/ml transforming growth factor beta one (TGFβ1, recombinant human, R&D systems) 25 μg/mL l-ascorbic acid 2-phosphate (Sigma Aldrich), 50 μg/ml gentamicin, and 1.5 μg/mL fungizone (both Gibco). After two weeks, TGFβ1 was no longer added to the medium and cells were cultured with and without 1μM of Forkosolin for 24h. Each experiment was performed with cells derived from 4 OA donors (3 Females, 1 Male, 65 ± 6 years), in triplicates.

### RNA isolation and real time quantitative PCR (RT-qPCR) for ATDC5 samples

Gene expression of *Col10a1* in ATDC5 expriments was evaluated by RT-PCR. For RNA isolation, chloroform was added to the TRIzol samples (TRIzol 5: Chlororform 1), which were subsequently centrifuged for 15min at 15000 rpm (i.e. RCF = 218849) and 4°C. RNA was isolated by collecting the aqueous phase and precipitated with isopropanol (aqueous phase 1: ispropanol 1) for 30min at −80°C. After centrifugation at 15000rpm (i.e. RCF = 218849) and 4°C for 30 min, supernatant was removed and the resulting pellet was washed with 80% Ethanol. RNA pellets were dried for 10min in desiccator and dissolved in 15μl RNase free water. Finally, RNA content and purity was determined with Nanodrop. RNA was converted to cDNA with the Revert Aid H Minus First strand cDNA synthesis kit (Thermo Scientific) according to the manufacturer’s protocols. Quantification of gene expression was done using Syber Select Master Mix (Applied Biosystems) adding 400nM forward and reverse oligonucleotides primers as listed in **supplementary method SM2.1**. The StepOne Plus System (Applied Biosystems) was used for amplification using the following protocol: denaturation cycle at 95°C for 10min followed by 40 cycles of amplification (15 seconds 95°C and 1 min 60°C), followed by a melting curve. Expression levels were analyzed using the 2^−ΔCt^ method and normalized for the expression of the reference gene *Hprt*. This housekeeping (HK) gene was determined after verification of multiple HK genes and selecting the one that remained most constant throughout the procedure.

The specific RNA isolation protocol, RT-qPCR method and oligonucleotides used for measuring gene expression in primary human chondrocytes is reported in **supplementary method SM2.2**.

### ALP assay

Enzymatic activity of secreted Alkaline Phosphatase (ALP) in the supernatant medium of ATDC5 cultures was determined in a colorimetric assay as previously described (*62*). Briefly, ALP activity was determined in flat-bottom 96-well plates (Sigma Aldrich, CAT M9410) containing assay buffer (1.5 M Tris-HCl, pH 9.0, 1 mM MgCl2; 7.5 mM p-nitrophenyl phosphate). The ALP activity was assessed as a function of formed nitrophenyl phosphate (pNp), the reaction colored product, which was measured by spectrophotometry at 405nM after 30min of reaction. The reaction was stopped with 1M NaOH Stop Buffer. A calibration curve containing an increasing concentration of pNp served to determine the absolute amount of ALP-generated pNp. Sample values were normalized to total DNA amount, expressed as μmol of pNp /mg of DNA and reported as fold change with respect to the relevant control medium.

### DNA quantification

After harvesting the cells with 350μl of 0.05% TritonX-100, samples were vortexed and frozen at −80°C for further processing. Samples were sonicated for 5 seconds, centrifuged for 1’ at 13000rpm (i.e. RCF= 164380) and 200μl of the supernatant was harvested. Samples were diluted with a factor 1/10 with distilled water then DNA content was measured with the Qubit 3.0 fluorometer (Life Technologies). Qubit dsDNA HS (high sensitivity, 0.2 to 500ng) Assay Kit was used according to the manufacturer’s protocols; a sample volume of 5μl was added to 195μl of a Qubit working solution.

### Statistical analysis

In general, one-tailed statistical tests were used to analyse *in vitro* results and relate them to expected outcomes from the *in silico* model’s predictions since the model predicts a directionality of the outcomes. Average effect of Forskoline and Activin treatments were compared to control in one representative experiment with 3 replicates thanks to a one-tailed unpaired t-test with welch’s correction, in **Fig5.C**. Effect of Forskoline treatment in OA chondrocytes from human donors was performed in triplicates for 4 donors. Comparision of the average effect with the control is done with a linear-mixed effect model to account for donor variability. Graphical visualization and statistical analyses for the semi-high throughput small molecule screening (**Fig.5D)** were performed by modifying the BraDiPluS package from the Saez Lab (https://saezlab.github.io/BraDiPluS/). More precisely, the probability that a treated condition resulted in lower ALP activity z-score than the corresponding control in the ATDC5 screening was estimated with a one-tailed Wilcoxon rank-sum test (the *in vitro* screening dataset was not normally distributed based on Shapiro-Wilk test), with Benjamini–Hochberg correction for multiple testing. Three independent experiments were performed for each conditions, and, treatments effects were assessed in triplicates, in each experiment (or run). Probabilities from the independent runs were combined with Fisher’s method using the combine.test function from the survcomp R package. A treated conditions was considered lower than control for p-value < 0.05 and z-score <0.

## 6. Data availability

### Code and data availability

The Matlab codes used to simulate and analyze the model as well as the R codes to analyze the experimental data are available on the GitHub page (https://github.com/Rapha-L/Insilico_chondro.git). The knowledge bases for the chondrocyte protein signaling (resp. gene regulatory) network are available through the Cellcollective online platform via this link (https://cellcollective.org/#a5b66073-6769-4c88-bf6b-37ca1aa8f766) (resp. this link: https://cellcollective.org/#474de240-8752-4c3b-aa63-23640e50bf7a).

## 7. Acknowledgements

## Fundings

This project has received funding from the European Union’s Horizon 2020 research and innovation programme under Marie Sklodowska-Curie grant agreement No 721432 and the European Research Council Consolidator Grant agreement No 772418, as well as the Fund for Scientific Research Flanders (FWO Vlaanderen) grant G085018N. The contribution of RN and GvO was provided within the framework of the Medical Delta Regmed4D program. The contribution of TW was supported by a grant from the Dutch Arthritis Association (LLP14).

The computational resources and services used in this work for parallel computing were provided by the VSC (Flemish Supercomputer Center), funded by the Research Foundation - Flanders (FWO) and the Flemish Government – department EWI.

The authors would like to thank D. Surtel, S. Ribeiro Viseu, O. Vanborren and J. Vleminckx for their technical support during the wet lab experiments.

## Supplementary Materials

**Supplementary computational method SM1**. Equations and mathematical framework.

**Supplementary experimental method SM2**. Oligonucleotide sequences and supplemental qPCR protocol.

**Fig. S1**. Individual dataset clustering and heatmap.

**Fig. S2.** Effect of ALKs ratio in the influence of inflammation and TGFβ signalings on chondrocyte hypertrophy.

**Fig. S3**. Screenshot of the user-friendly interface for the virtual chondrocytes App.

**Fig. S4.** Decision tree summarizing the variable updating scheme employed in algorithm to simulate the in silico chondrocyte

**Fig. S5**. Illustration of the algorithm for the asynchronous updating of variables with a simplified (3 nodes) example network.

**Table S1**. Definition box.

**Table S2**. List of variables and mouse gene correspondence.

**Data S1**. Subset of genes for gene expression profiles and description of GEO datasets with manual binary annotations.

**Data S2**. Attractors complete protein and gene expression profiles.

**Data S3.** In silico screening predictions of combinatorial treatments.

