## Supplementary material for "An integrated *in silico*-*in vitro* approach for identification of therapeutic drug targets for osteoarthritis"

### Supplementary Materials for

#### An integrated in silico-in vitro approach for the identification of drug targets against chondrocyte hypertrophy in osteoarthritis

Raphaëlle Lesage, Mauricio N. F. Blanco, Roberto Narcisi, Tim Welting, Gerjo J.V.M. Van Osch, Liesbet Geris*

**This PDF file includes:**

Supplementary Text
Figs. S1 to S5
Tables S1 to S2
References (1)

**Other Supplementary Materials for this manuscript include the following:**

Data S1 to S3

**Supplementary Text**

Mathematical framework and system of equations for the articular chondrocyte regulatory network:

- *Theoretical frame:*

The potential function of a component (or node) on its downstream neighbours is called the global activity. It corresponds to the multiplication of the gene activation level by the protein activation level.

Let's consider the adjacency matrix $A$ of the network whose entries $a_{ij}$ can be 0, 1 or -1. This matrix indicates the presence and the direction of edges in the network. If node $j$ activates node $i$, then $a_{ij}$ is 1; if node $j$ inhibits node $i$, $a_{ij}$ is -1. $a_{ij}$ equals 0 when no interaction from node $j$ to node $i$ is present. The vector $L = \ l_{ij}$ contains a weight for each interaction in the regulatory network. Given $A$ and $L$, the full system of equations contains equations for both the fast and the slow reactions regulating each variable and can be written as:

$$\begin{cases} z_1^f(t+1) = a_{11}^f l_{11}^f\, z_1(t) + a_{12}^f l_{12}^f\, z_2\,(t) \ldots + a_{1n}^f l_{1n}^f\, z_n(t) \\ \ldots \\ z_n^f(t+1) = a_{n1}^f l_{n1}^f\, z_1(t) + a_{n2}^f l_n^f\, z_2\,(t) \ldots + a_{nn}^f l_{nn}^f\, z_n(t) \\ z_1^s(t+1) = a_{11}^s l_{11}^s\, z_1(t)\ + a_{12}^s l_{12}^s\, z_2\,(t) \ldots + a_{1n}^s l_{1n}^s\, z_n(t) \\ \ldots \\ z_n^s(t+1) = a_{n1}^s l_{n1}^s\, z_1(t) + a_{n2}^s l_n^s z_2\,(t) \ldots + a_{nn}^s l_{nn}^s\, z_n(t) \end{cases} \qquad \text{(eq.1)}$$

Or

$$\begin{cases} \boldsymbol{z}^f(t+1) = A^f L^f \boldsymbol{z}(t) \\ \boldsymbol{z}^s(t+1) = A^s L^s \boldsymbol{z}(t) \end{cases} \qquad \text{(eq.2)}$$

and the global activity of the component $i$ at the time $t+1$ is defined as

$$\boldsymbol{z_i(t+1) = z_i^f(t+1) \times z_i^s(t+1)} \qquad \text{(eq.3)}$$

where $n$ is the total number of nodes in the network, **z** is a vector containing the activities for all nodes, $A^v = [a_{ij}^v]$ and $L^v = \left[l_{ij}^v\right]$are matrices with $v \in \{s, f\}$ and $\mathrm{i,j} \in [1, n]$). $f$ and $s$ denote fast and slow variables, respectively. $z^f$ and $z^s$ and $\boldsymbol{z}$ are $n \times 1$ vectors filled with the $z_i^f, z_i^s\ or\ z_i$ elements respectively. When two nodes are known to act in complex and in synergy, their individual terms are merged. Indeed, a term $a_{i(j,k)} l_{i(j,k)} z_j z_k$ replaces $a_{ij} l_{ij} z_j + a_{ik} l_{ik} z_k$ in the equations (**eq.1**).

The additive sum includes a weight ($l$) for each term that assumes a limited amount of saturation whenever there is a majority of stimulatory interactions. The value of that saturation depends on the interactions' signs as defined by Kerkhofs et al. in (*1*):

$$l_{ij} = \begin{cases} 1, & if \sum_{j \in U} a_{ij} \leq 1 \\ 2 \dfrac{SC}{\sum_{j \in U} a_{ij}}, & if \sum_{j \in U} a_{ij} > 1 \end{cases} \qquad \text{(eq.4)}$$

where $U$ is the set of nodes' indexes that are upstream node $i$ and $SC$ is the saturation constant. If the positive interactions do not outnumber the inhibitory interactions by more than 1, all weights are set to 1. If the number of excess positive interactions is higher than 1, we introduce a saturation factor whose value will determine how fast the node will saturate. The saturation constant was set to $\frac{2}{3}$ in that study based on (*1*). The table below (**Table sup. text 1**) shows the weights for various values of $\sum_{j \in U} a_{ij}$ (the number of excess positive interactions). We refer the reader to the paper of Kerkhofs et al. (*1*) for more information pertaining to the implication of the saturation term and sensitivity to the saturation constant value.

| $\sum_{j \in U} a_{ij}$ (excess positive interactions) | 2 | 3 | 4 | 5 |
|---|---|---|---|---|
| $l_{ij}$ (weight) | $\frac{2}{3} \approx 0.66$ | $\frac{4}{9} \approx 0.44$ | $\frac{1}{3} \approx 0.33$ | $\frac{4}{15} \approx 0.267$ |

**Table sup. text 1**: Weight for various values of the number of excess positive interactions and for the saturation constant set to 2/3.

Overall, while these rules cover a standard situation, incorporation of relevant biological facts should always prevail over these a priori rules. For that reason, some adaptations to the weights in the chondrocyte network were introduced to better reflect natural expression profiles in the WT situation.

Importantly, this additive model was semi-quantitative since the nodes or variables could take on a continuous activity value between 0 and 1. For a given sub-variable (i.e. fast or slow) if the sum and substraction of activator and inhibitory influences was higher than 1 (respectively lower than 0), then the component was considered as fully activated (resp. inhibited) and the value was brought back to exactly 1 (resp. 0). In mathematical terms:

$$z^{v}(t) = \begin{cases} 0, & if\ z^{v}(t) \leq 0 \\ 1, & if\ z^{v}(t) \geq 1 \end{cases} \quad \text{(eq.5)}$$

with $v \in \{s, f\}$.

- *Application to the chondrocyte network:*

In the articular chondrocyte regulatory network (c.f. **Fig. 1**), the global activity of a variable $i$ is defined as follow (c.f. **eq.3**) :

$$\boldsymbol{z_i(t) = z_i^f(t) \times z_i^s(t)} \quad \text{(eq.6)}$$

with $i \in [1, 60]$, indicating the variable number.

The full system of reaction is provided in two parts: (a) the fast reactions, (b) the slow reactions, the variable index refers to the index provided in **Table S2**.

a) <u>Fast reactions (protein signaling network)</u>

$$z_1^f(t+1) = 1 - z_{49}(t) - z_{48}(t)$$
$$z_2^f(t+1) = z_{50}(t) - 0.3 * z_4(t)$$
$$z_3^f(t+1) = 1$$
$$z_4^f(t+1) = z_{52}(t) - 0.5 * z_{25}(t) + z_{52}(t)$$

$$z_5^f(t+1) = 1$$
$$z_6^f(t+1) = \frac{2}{3} * z_5(t)$$
$$z_7^f(t+1) = 1 - z_{36}(t)$$
$$z_8^f(t+1) = z_7(t) - z_{10}(t)$$
$$z_9^f(t+1) = z_{38}(t) + z_{22}(t) + z_{32}(t) - z_{10}(t) - z_{14}(t) - z_{26}(t) \times z_{30}(t) - z_{31}(t) - z_{43}(t)$$
$$z_{10}^f(t+1) = z_{14}(t) + z_{26}(t) - z_7(t) - z_{19}(t)$$
$$z_{11}^f(t+1) = 1$$
$$z_{12}^f(t+1) = z_{11}(t)$$
$$z_{13}^f(t+1) = 1$$
$$z_{14}^f(t+1) = \left(z_{12}(t) + z_3(t)\right) \times s_2$$
$$z_{15}^f(t+1) = z_{34}(t) + z_{19}(t) - z_{30}(t) - z_{26}(t)$$
$$z_{16}^f(t+1) = 1$$
$$z_{17}^f(t+1) = z_{16}(t)$$
$$z_{18}^f(t+1) = \frac{2}{3} \times \left(\frac{2}{3} \times z_{17}(t) + \frac{2}{3} \times z_{27}(t)\right) + \frac{1}{3} \times z_{42}(t) - z_{19}(t) + z_{54}(t) \times (1 - z_{17}(t))$$
$$z_{19}^f(t+1) = z_4(t) - 0.25 \times z_{22}(t)$$
$$z_{20}^f(t+1) = 1$$
$$z_{21}^f(t+1) = 1$$
$$z_{22}^f(t+1) = z_{41}(t) - z_{37}(t) + z_{55}(t)$$
$$z_{23}^f(t+1) = 1$$
$$z_{24}^f(t+1) = 1$$
$$z_{25}^f(t+1) = 1$$
$$z_{26}^f(t+1) = z_{53}(t) - 0.5 \times \left(\left(z_{22}(t) + z_{31}(t) + z_{25}(t)\right) \times s_3\right)$$
$$z_{27}^f(t+1) = z_{16}(t)$$
$$z_{28}^f(t+1) = \left(z_{34}(t) + z_{34}(t) \times z_{26}(t)\right) \times s_2$$
$$z_{29}^f(t+1) = \left(z_{27}(t) + z_{38}(t) + z_{55}(t) - z_{58}(t)\right) \times s_2$$
$$z_{30}^f(t+1) = z_{37}(t) - z_{35}(t)$$
$$z_{31}^f(t+1) = 1 - 0.5 \times \left(z_{35}(t) + z_{18}(t)\right)$$
$$z_{32}^f(t+1) = z_{34}(t) + z_{19}(t) - z_{43}(t)$$
$$z_{33}^f(t+1) = 1 - z_{47}(t)$$
$$z_{34}^f(t+1) = \left(z_{23}(t) + z_{33}(t) + z_{55}(t)\right) \times s_3$$
$$z_{35}^f(t+1) = 1 - z_{14}(t) - z_{38}(t)$$
$$z_{36}^f(t+1) = 1 + z_{37}(t) - z_2(t) - 0.5 \times z_{22}(t)$$
$$z_{37}^f(t+1) = z_{14}(t)$$
$$z_{38}^f(t+1) = z_{39}(t) - 0.5 \times z_{37}(t)$$
$$z_{39}^f(t+1) = \left(z_{41}(t) + z_{42}(t) + z_{54}(t)\right) \times s_3$$

$$z_{40}^{f}(t+1) = \left(z_{22}(t) + z_{56}(t)\right) \times s_2$$
$$z_{41}^{f}(t+1) = \left(z_{1}(t) + z_{33}(t) + z_{17}(t) + z_{27}(t)\right) \times s_4$$
$$z_{42}^{f}(t+1) = z_{3}(t)$$
$$z_{43}^{f}(t+1) = 1 - z_{32}(t)$$
$$z_{44}^{f}(t+1) = 1$$
$$z_{45}^{f}(t+1) = \left(z_{14}(t) + z_{22}(t)\right) \times s_2$$
$$z_{46}^{f}(t+1) = 1$$
$$z_{47}^{f}(t+1) = 1$$
$$z_{48}^{f}(t+1) = 1$$
$$z_{49}^{f}(t+1) = 1$$
$$z_{50}^{f}(t+1) = z_{1}(t) - z_{49}(t)$$
$$z_{51}^{f}(t+1) = 1$$
$$z_{52}^{f}(t+1) = z_{23}(t) - z_{33}(t)$$
$$z_{53}^{f}(t+1) = z_{23}(t)$$
$$z_{54}^{f}(t+1) = z_{51}(t) - z_{59}(t)$$
$$z_{55}^{f}(t+1) = z_{54}(t) - 0.5 \times z_{34}(t)$$
$$z_{56}^{f}(t+1) = z_{55}(t)$$
$$z_{57}^{f}(t+1) = 1 - z_{24}(t)$$
$$z_{58}^{f}(t+1) = 1 - z_{55}(t)$$
$$z_{59}^{f}(t+1) = 1$$
$$z_{60}^{f}(t+1) = 0.75 + z_{56}(t) - z_{38}(t) - z_{22}(t)$$

b) Slow reactions (gene regulatory network)

$$z_{1}^{s}(t+1) = 2 \times z_{6}(t) - z_{43}(t)$$
$$z_{2}^{s}(t+1) = 1$$
$$z_{3}^{s}(t+1) = z_{14}(t)$$
$$z_{4}^{s}(t+1) = 1 - z_{45}(t) + z_{44}(t)$$
$$z_{5}^{s}(t+1) = \left(z_{9}(t) + z_{19}(t) + z_{29}(t) + z_{45}(t) - z_{44}(t) - z_{17}(t)\right) \times s_2$$
$$z_{6}^{s}(t+1) = 1 - z_{17}(t)$$
$$z_{7}^{s}(t+1) = 1$$
$$z_{8}^{s}(t+1) = \left(1 + z_{9}(t) + z_{7}(t)\right) \times s_3$$
$$z_{9}^{s}(t+1) = \left(z_{46}(t) + z_{8}(t) + z_{9}(t) + z_{15}(t) + z_{32}(t) - z_{21}(t) \times z_{19}(t) - z_{43}(t) + z_{6}(t) - z_{14}(t) + z_{40}(t)\right) \times s_4$$
$$z_{10}^{s}(t+1) = \left(z_{14}(t) - z_{29}(t) + z_{34}(t) + z_{29}(t) + z_{10}(t) + z_{21}(t)\right) \times s_4$$
$$z_{11}^{s}(t+1) = \left(z_{6}(t) \times z_{10}(t) + z_{10}(t) + z_{26}(t)\right) \times s_3$$
$$z_{12}^{s}(t+1) = \left(z_{6}(t) + z_{10}(t) + z_{19} - z_{18}(t)\right) \times s_2$$
$$z_{13}^{s}(t+1) = \left(z_{9}(t) + z_{4}(t) + z_{15}(t) - z_{14}(t) - z_{11}(t) + z_{46}(t) - z_{18}(t) + z_{7}(t)\right) \times s_2$$
$$z_{14}^{s}(t+1) = 1$$
$$z_{15}^{s}(t+1) = \left(z_{9}(t) + z_{19}(t)\right) \times s_2$$

$$z_{16}^{s}(t+1) = \left(z_7(t) + z_9(t)\right) \times s_2$$
$$z_{17}^{s}(t+1) = z_{10}(t) - z_{40}(t) \times z_{22}(t)$$
$$z_{18}^{s}(t+1) = 1$$
$$z_{19}^{s}(t+1) = 1$$
$$z_{20}^{s}(t+1) = \left(z_{10}(t) + z_{27}(t) + z_{21}(t) - z_7(t) + z_{60}(t)\right) \times s_3$$
$$z_{21}^{s}(t+1) = \left(z_{10}(t) + z_{14}(t)\right) \times s_2$$
$$z_{22}^{s}(t+1) = z_{15}(t)$$
$$z_{23}^{s}(t+1) = z_6(t) - z_{40}(t) + z_{26}(t)$$
$$z_{24}^{s}(t+1) = z_9(t) + z_{29}(t) + z_{46}(t) + z_6(t) + z_{46}(t) \times z_9(t) \times z_{34}(t) + z_{19}(t) + z_{40}(t) - z_{26}(t) + z_{40}(t) \times z_{28}(t) \times z_{56}(t) - \frac{1}{5} z_{60}(t)$$
$$z_{25}^{s}(t+1) = \left(z_{18}(t) + z_{19}(t) + z_{26}(t) + 2 \times z_{29}(t)\right) \times s_5$$
$$z_{26}^{s}(t+1) = 1$$
$$z_{27}^{s}(t+1) = z_9(t) - 0.5 \times z_{19}(t)$$
$$z_{29}^{s}(t+1) = 1$$
$$z_{30}^{s}(t+1) = 1$$
$$z_{31}^{s}(t+1) = \left(z_{28}(t) + z_6(t) + 1.5 \times z_{14}(t) + z_{60}(t) \times z_{26}(t)\right) \times s_4$$
$$z_{32}^{s}(t+1) = z_{15}(t) + z_{19}(t) - z_{26}(t)$$
$$z_{33}^{s}(t+1) = \left(z_6(t) + z_{29}(t)\right) \times s_2$$
$$z_{34}^{s}(t+1) = 1 - 0.4 \times z_{44}(t)$$
$$z_{35}^{s}(t+1) = 1$$
$$z_{36}^{s}(t+1) = 1$$
$$z_{37}^{s}(t+1) = \left(1 - z_7(t)\right) \times s_2$$
$$z_{38}^{s}(t+1) = z_9(t)$$
$$z_{39}^{s}(t+1) = z_9(t)$$
$$z_{40}^{s}(t+1) = 1$$
$$z_{41}^{s}(t+1) = 1$$
$$z_{42}^{s}(t+1) = \left(z_{26}(t) + z_{29}(t) + z_{18}(t)\right) \times s_3$$
$$z_{43}^{s}(t+1) = \left(z_{19}(t) + z_{26}(t) + z_{43}(t) - z_{32}(t)\right) \times s_2$$
$$z_{44}^{s}(t+1) = \left(z_{26}(t) + z_{29}(t) + z_{40}(t) - z_{19}(t) + z_4(t)\right) \times s_3$$
$$z_{45}^{s}(t+1) = 1$$
$$z_{46}^{s}(t+1) = z_{29}(t)$$
$$z_{47}^{s}(t+1) = z_{10}(t) - z_7(t)$$
$$z_{48}^{s}(t+1) = 0.5 \times z_{19}(t) - z_7(t) + z_{10}(t) - z_{29}(t) - z_{28}(t)$$
$$z_{49}^{s}(t+1) = 0.5 \times z_{19}(t) - z_7(t) + z_{10}(t) - z_{29}(t) - z_{28}(t)$$
$$z_{50}^{s}(t+1) = 1$$
$$z_{51}^{s}(t+1) = z_{40}(t) \times z_{28}(t) \times z_{56}(t) + z_{29}(t)$$
$$z_{52}^{s}(t+1) = \left(0.4 + z_{18}(t) - z_{26}(t) + z_9(t) + z_{28}(t) \times z_{56}(t)\right) \times s_3$$
$$z_{53}^{s}(t+1) = \left(0.4 + z_{26}(t) - z_{29}(t) + z_{10}(t)\right) \times s_2$$
$$z_{54}^{s}(t+1) = 1 - z_{60}(t)$$
$$z_{55}^{s}(t+1) = 1$$
$$z_{56}^{s}(t+1) = 1$$
$$z_{57}^{s}(t+1) = \left(z_{10}(t) + z_{60}(t) - z_{29}(t)\right) \times s_3$$

$$z_{58}^{s}(t+1) = z_{29}(t)$$
$$z_{59}^{s}(t+1) = \left(z_{42}(t) + z_{17}(t)\right) \times s_3$$
$$z_{60}^{s}(t+1) = z_{26}(t) - z_{40}(t) \times z_{28}(t) - z_{29}(t)$$

With $s_p = 2^{\frac{SC}{p}}$, where *SC* is the saturation constant set to 2/3 and *p* is the number of excess positive interactions (see equation **eq.4**).

Supplementary experimental method and sequences of primers for qPCR:

**Mouse oligonucleotides used for qRT-PCR reaction with ATDC5 samples.** Primers were designed by the Primer Design tool of NCBI:

| Gene | Forward primer | Reverse primer |
|---|---|---|
| m*Hrpt* | GAGCGTTGGGCTTACCTCAC | ATCGTAATCACGACGCTGG |
| m*Col10a1* | TCCCAGCACCAGAATCTATCTGA | TTATGCCTGTGGGCGTTTGG |

**RT-qPCR protocol for samples from human primary OA chondrocytes cultured in alginate beads:** Alginate beads were dissolved using citrate buffer, centrifuged at 200g and the pellet was resuspended in RLT (Qiagen, Hilden, Germany) buffer containing 1% beta-mercaptoethanol for RNA isolation. mRNA isolation was performed according to manufacturer's protocol utilizing the RNeasy Column system (Qiagen, Hilden, Germany). The RNA concentration was determined using a NanoDrop spectrophotometer (Isogen Life Science, Utrecht, the Netherlands). 0.5 µg RNA was used for cDNA synthesis following the protocol of the manufacturer of the RevertAid First Strand cDNA kit (Thermo Fisher Scientific, Waltham, MA, United States). qPCR was performed on a Bio-Rad CFX96 Real-Time PCR Detection System (Bio-Rad) to assess gene expression, Collagen type 10 (*COL10A1*) and Glyceraldehyde-3-phosphate dehydrogenase (*GAPDH*), which was found stable and therefore used as reference gene. Data were analyzed by the ΔΔCt method and normalized to the expression of *GAPDH* of each condition and compared to the corresponding gene expression in the control groups

**Human oligonucleotides used for qRT-PCR reaction with primary chondrocytes:**

| Gene | Forward primer | Reverse primer | Probe |
|---|---|---|---|
| *GAPDH* | ATGGGGAAGGTGAAGGTCG | TAAAAGCAGCCCTGGTGACC | CGCCCAATACGACCA AATCCGTTGAC |
| *COL10A1* | CAAGGCACCATCTCCAGGAA | AAAGGGTATTTGTGGCAGCATATT | TCCAGCACGCAGAAT CCATCTGA |

GSE26475

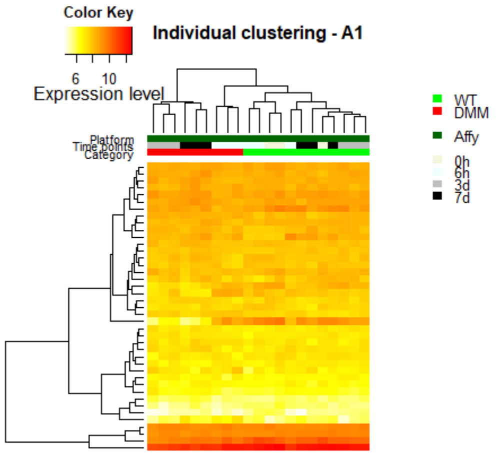


GSE33754

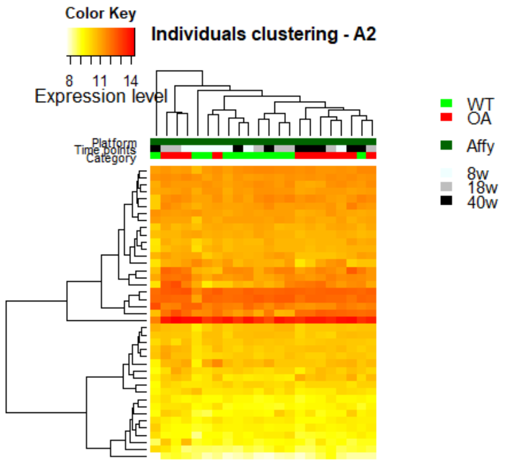


GSE33656

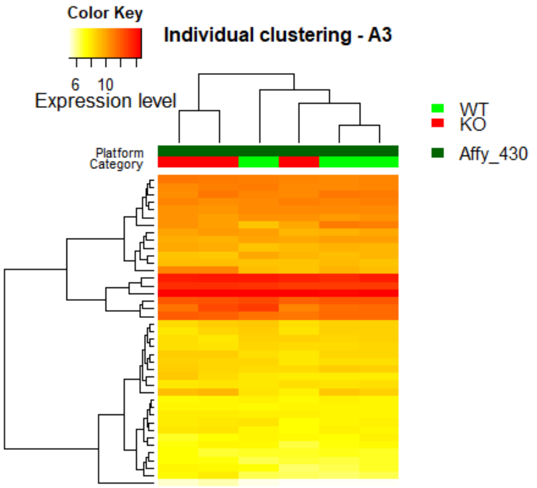


GSE45793

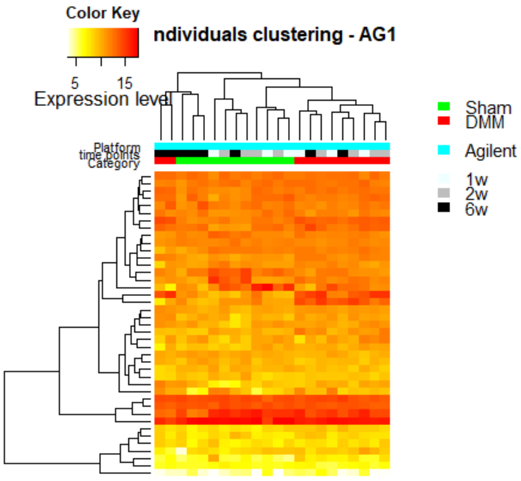


GSE79239

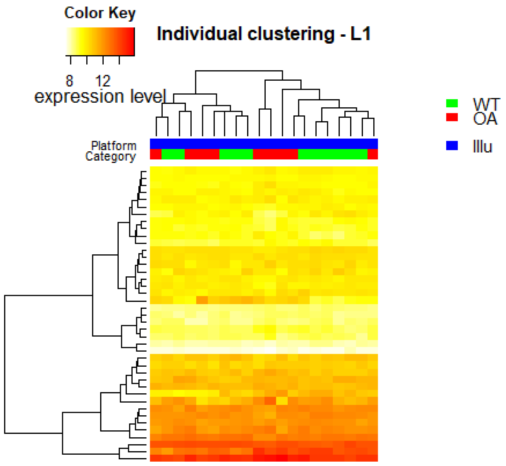


GSE53857

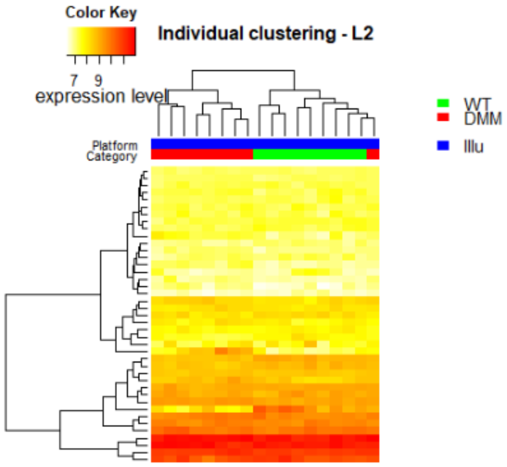

**Fig. S1. Heatmap with unsupervised clustering on individual dataset.**

The microarray sub-datasets, used in the merged dataset, were also investigated individually. They were pre-processed (i.e. processing steps before merging and correcting for batch effect) in the same way as the merged dataset but the unsupervised clustering analysis was done on each sub-dataset separately in order to compare the biological or OA related information content before and after the data merging. The heatmaps show the expression profile of the same list of genes of interest than for the merged dataset (see **Data S3**.). These expression datasets were submitted to unsupervised clustering with the Euclidean distance method and the Complete aggregation method in R thanks to the heatmap3 function from the github repository https://github.com/obigriffith/biostar-tutorials/tree/master/Heatmaps. The headers indicate the GEO accession numbers of the 6 original datasets. Samples labeled as 'WT', for wild type, are in green, samples labeled as 'OA', for osteoarthritis, in red. The pre-labelling is the same as for the merged dataset (see **Data S3**). When applicable, a grey scale indicate the time points (w stands for weeks, in the legend). For some datasets (e.g. GSE26475, GSE33656, GSE53857) the OA and the WT samples are well separated in different clusters, while for other the separation is not so clear. For instance, in GSE45793 the variance due the time point 6weeks is greater than the OA induced variance, while for weeks 1 and 2 OA and WT samples are well separated due to the OA condition.

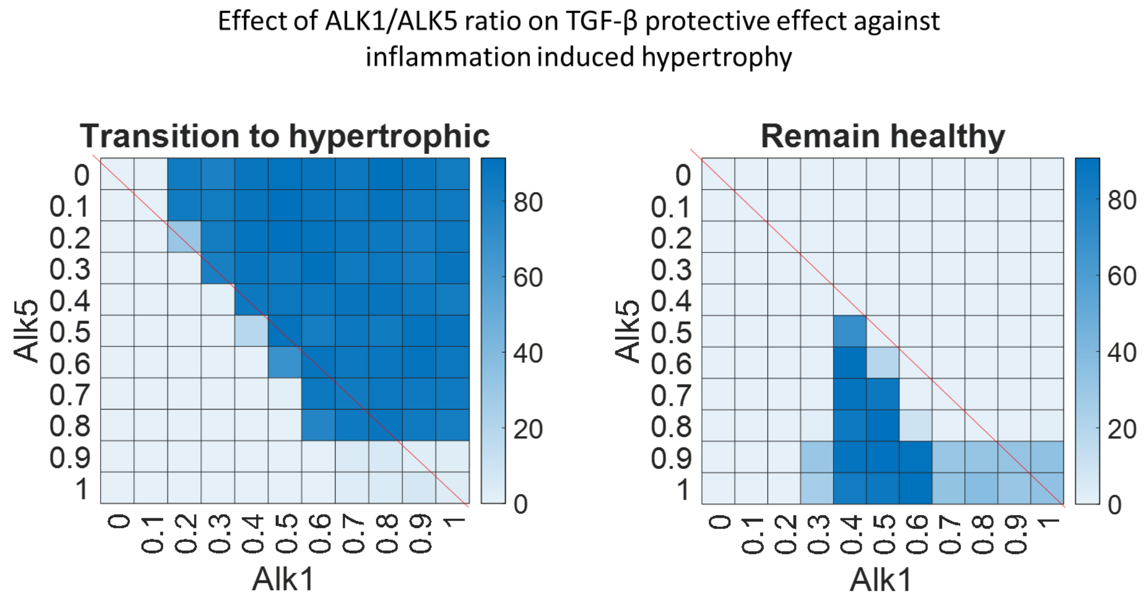


Fig.

**Fig S2. Role of the ratio between receptors ALK1 and ALK5 in the effect of inflammation and TGFβ signaling on chondrocyte hypertrophy**

Percentage of perturbations remaining in the healthy state (right) or transitioning towards the hypertrophic one (left) during inflammatory pathway activation with TGF-B treatment while changing the ratio between ALK1 and ALK5. The inflammatory and TGF-B profiles that were imposed are the same as in Fig.4A except that the value imposed for ALK1 and ALK5 are varying between 0 and 1 with a 0.1 increment. ALK1=ALK5 on the red diagonals and ALK1>ALK5 in the upper right corner. Roughly, the rescue of the healthy state by TGF-B is lost when ALK1 is greater than ALK5. If ALK1 is high enough and that the difference between ALK1 and ALK5 is not greater than 10-20% then the protective effect of TGF-B is also disturbed for ALK1<ALK5

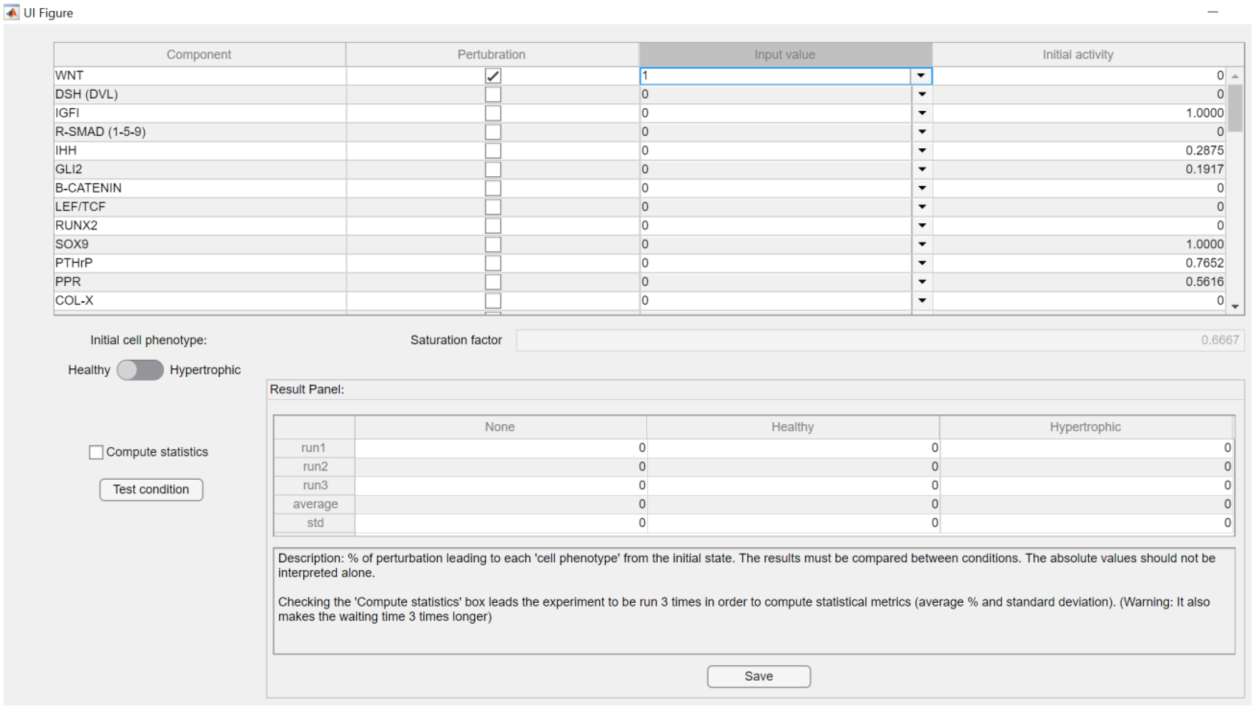


**Fig. S3. Screenshot of the user-friendly interface for the virtual chondrocytes App**

The standalone Matlab-based applications can be launched and used without Matlab license, provided that the compiler Matlab Runtime is installed (https://nl.mathworks.com/products/compiler/matlab-runtime.html ). The virtual chondrocyte initial state can be set as healthy or hypertrophic, allowing the user to test any scenarios. All the 60 components may be perturbed alone or in any sort of combination by forcing the variables to take a value in the interval [0:1], with a step of 0.1. The most left column indicate the value of the variable in the selected initial state, for information. Obviously, applying a perturbation that is equal to the initial value of the variable will not affect the system. Once the setting are done, the user can apply the experimental condition by pushing the button ‘Test condition’ and the percentage of transition towards each of the possible basal stable states (i.e. ‘None’, ‘Healthy’ and ‘Hypertrophic’) is computed. If the ‘Compute statistics’ box is ticked, then the experiment is repeated 3 times and the average and standard deviations are displayed (variation occurs due to the stochastic nature of the model). The results may be exported and saved in an excel file via the ‘Save’ button.

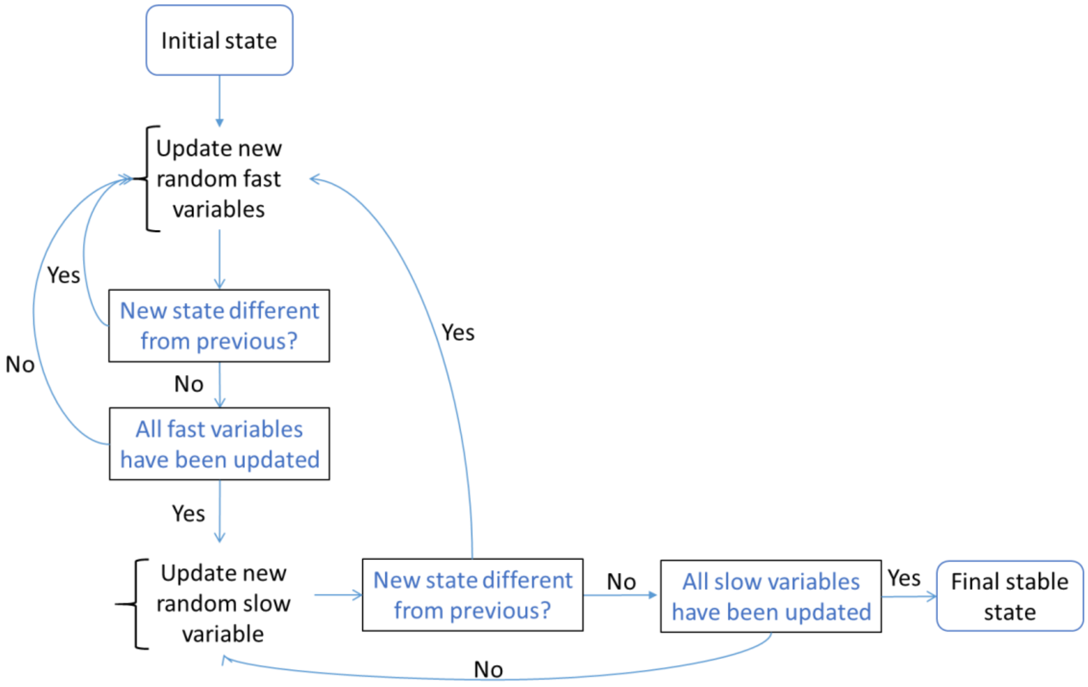


**Fig. S4. Decision tree summarizing the variable updating scheme employed in algorithm to simulate the *in silico* chondrocyte**

Each biological component is represented by the gene expression level (slow variable) and the protein activity potential (fast variable). Variables are updated based on the rules stored in the model's equations. First, fast variables are updated in random order, when a pseudo-stable state is reached and that all fast variables have been updated, the next random chosen slow variable is updated. This goes on until a state that is stable both at the fast and slow level is reached. This is the final stable state. A state is considered stable if further variable updates do not bring further changes for any of the variables, more or less a predefined tolerance. The order in which variables are updated is random, thereby generating some stochasticity in the model. Within the fast (resp. slow) updating loops, variables are updated asynchronously (meaning the one after the others) according to the rules defined in the system of equations (see Supplementary text S1) and in a random order.

The asynchronous updating system is illustrated for a simplified example network below:

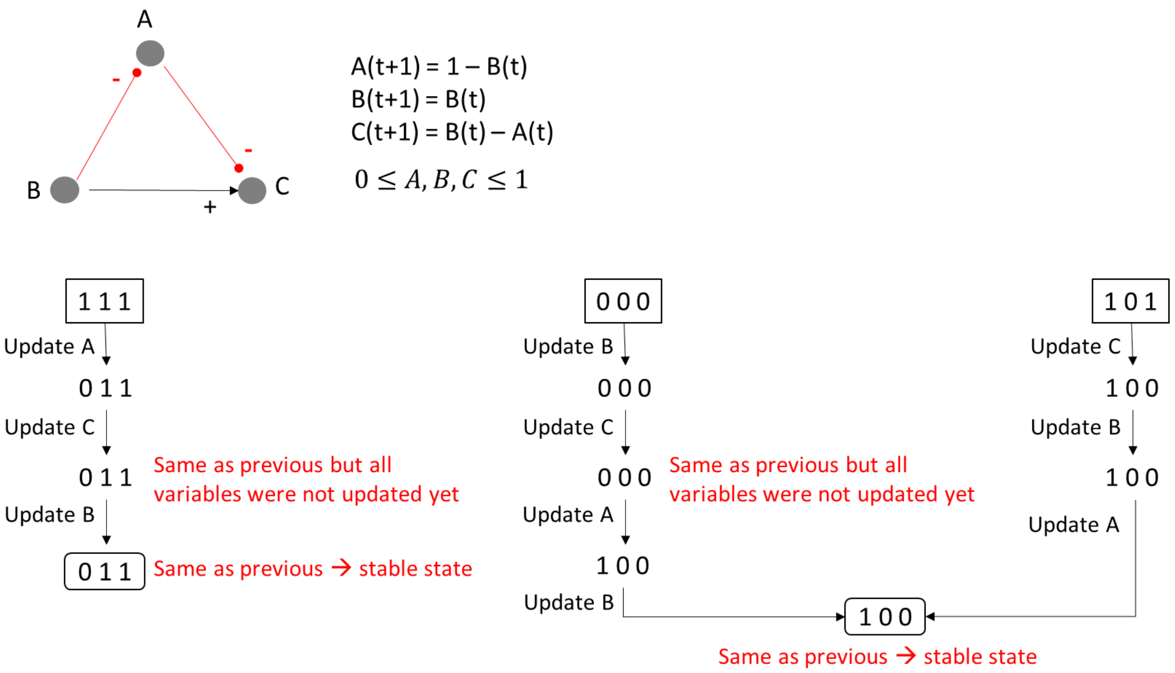


**Fig. S5. Illustration of the algorithm for the asynchronous updating of variables with a simplified example network**

The network represents interactions happening in one of the layers (protein= fast reactions or genetic = slow reactions). Inhibitory influences are represented in Red and activator influences are in black. The mathematical rules corresponding to the network are displayed. If the rules result in a value lower than 0 (resp. higher than 1), the value is brought back to 0 (resp. 1). For this example, 3 different initial states are inputted and each of the three variables is updated asynchronously. The order in which variables are updated is random. The system reaches a stable state when the next update gives the same state as in the previous time step and that all variables were screened in the random ordering list. That state is a pseudo-stable state if the rules were describing fast reactions, in that case, a new slow variable can be updated (see **Fig. S3**.). It is a final stable state if the rules were describing slow interactions since it would mean that the system had first reached a pseudo-stable state at the fast level and would now be stable at the slow level too.

**Table S1. Table of definitions.** Modelling terms and concepts relative to the current additive regulatory network model are defined.

| Definitions |
|---|
| **Additive model**: A network of protein or gene interactions is modeled with an additive method if the evolution of variables (proteins or genes) at the next time step is defined by the sum of the upstream activating variables and the subtraction of the upstream inhibitory variables of the network. For instance, in a GRN, if a transcription factor A activates a gene P while B inhibits gene P, then the evolution of the P expression level at the next time step is defined as: $\mathrm{P(t+1)} = \mathrm{A(t)} - \mathrm{B(t)}$ (eq. 1) |
| **Node or component (variable):** The nodes of a regulatory network are located at the intersections of multiple interactions (edges) in the network. They represent biological components such as proteins or genes. In mathematical models, such as the additive models, components' evolutions are described with variables (e.g. $A$, $P$ or $B$ in (eq.1) ). |
| **Fast and Slow reactions & variables:** All reactions related to slow biological processes such as gene expression, mRNA or protein production, were referred to as slow reactions (lower priority) and those related to fast processes such as protein activation (e.g. post transcriptional modification) or degradation, were referred to as fast reactions (higher priority).<br>This priority order plays a role in the simulation since any network component could be regulated both at the protein and gene level, as in real life. Therefore, each network component or variable was split into a fast and a slow subpart, also called sub-variables. Let's consider that the gene P from (eq.1) produces a protein that is both activated post-transcriptionally by another protein kinase K and blocked by an inhibitory protein I. The global functional activity of the protein P is actually the multiplication of the slow by the fast subparts:<br>$\mathrm{P(t+1)} = [\,\mathrm{A(t)} - \mathrm{B(t)}\,] \times [\,\mathrm{K(t)} - \mathrm{I(t)}\,]$. (eq2)<br>When simulating the system, the sub-variables are updated asynchronously following the priority classes in such a way that fast reactions are always updated before the slow reactions (*20*). |
| **Stable state (or attractors):** A stable state is an ensemble of values, one for each variable/component of the model, that meet all the rules/constraints imposed by the equations. When variables do not evolve anymore after a certain time of simulation, the system has reached a stable state. The nature of such a state depends on the system of equations and on which initial state was used. Given the ensemble of interconnected signaling pathways, it is likely that only a finite number of states can fulfil all the constraints imposed by the network structure (i.e. by the equations). |

**Table S2. Table of correspondence between the variable names in the model corresponding real world mouse gene.** All mathematical variables and the corresponding node in the network have names written in upper cases and do not reflect the official human or mouse nomenclature. To relate those variables to actual genes more easily, we provide this table of correspondence. A related mouse gene name and NCBI ID is indicated for each variable. Nevertheless, it is not exhaustive since some variables represents a group of factors or a family of ligands rather than a single factor.

| Variable index | Variable name | Example representative mouse gene | |
|---|---|---|---|
| | | Name | NCBI ID |
| 1 | WNT | *Wnt3a* | 22416 |
| 2 | DSH | *Dvl1* | 20423 |
| 3 | IGF-I | *Igf1* | 16000 |
| 4 | R-SMAD | *Smad5* | 17129 |
| 5 | IHH | *Ihh* | 16147 |
| 6 | GLI2 | *Gli2* | 14633 |
| 7 | β-Catenin | *Ctnnb1* | 12387 |
| 8 | LEF/TCF | *Tcf7* | 21414 |
| 9 | RUNX2 | *Runx2 or Cbfa1* | 12393 |
| 10 | SOX9 | *Sox9* | 20682 |
| 11 | PTHrP | *Pthlh* | 19227 |
| 12 | PPR | *Pth1r* | 19228 |
| 13 | COL-X | *Col10a1* | 12813 |
| 14 | PKA | *Prkaca* | 18747 |
| 15 | MEF2C | *Mef2c* | 17260 |
| 16 | FGF | *Fgf2* | 14173 |
| 17 | FGFR3 | *Fgfr3* | 14184 |
| 18 | STAT1 | *Stat1* | 20846 |
| 19 | Smadcomplex | *Smad4 & R-Smads* | 17128 |
| 20 | COL II | *Col2a1* | 12824 |
| 21 | NKX3.2 | *Nkx3-2 or Bapx1* | 12020 |
| 22 | ERK1/2 | *Mapk3* | 26417 |
| 23 | TGFβ | *Tgfb1* | 21803 |
| 24 | MMP13 | *Mmp13* | 17386 |
| 25 | SMAD7 | *Smad7* | 17131 |
| 26 | SMAD3 | *Smad3* | 17127 |
| 27 | FGFR1 | *Fgfr1* | 14182 |
| 28 | ATF2 | *Atf2* | 11909 |
| 29 | NFκB | *Nfkb1* | 18033 |

| | | | |
|---|---|---|---|
| 30 | HDAC4 | *Hdac4* | 208727 |
| 31 | CCND1 | *Ccnd1* | 12443 |
| 32 | DLX5 | *Dlx5* | 13395 |
| 33 | BMP | *Bmp2* | 12156 |
| 34 | P38 | *Mapk14* | 26416 |
| 35 | GSK3β | *Gsk3b* | 56637 |
| 36 | DC | *Apc* | 11789 |
| 37 | PP2A | *Ppp2ca* | 19052 |
| 38 | AKT | *Akt1* | 11651 |
| 39 | PI3K | *Pi3kr1* | 18708 |
| 40 | ETS1 | *Ets1* | 23871 |
| 41 | RAS | *Kras* | 16653 |
| 42 | IGF-IR | *Igf1r* | 16001 |
| 43 | MSX2 | *Msx2* | 17702 |
| 44 | δEF-1 | *Zeb1* | 21417 |
| 45 | ATF4 | *Atf4 or Creb* | 11911 |
| 46 | HIF-2α | *Epas1* | 13819 |
| 47 | GREM1 | *Grem1* | 23892 |
| 48 | DKK1 | *Dkk1* | 13380 |
| 49 | FRZB | *Frzb* | 20378 |
| 50 | Frizzled-LRP5/7 | *Lrp5* | 16973 |
| 51 | Cytokines | *Il1b* | 16176 |
| 52 | ALK1 | *Acvrl1* | 11482 |
| 53 | ALK5 | *Tgrb1* | 21812 |
| 54 | R-Infl (Receptor inflammation) | *Tlr1* | 21897 |
| 55 | TAK1 | *Map3k7* | 26409 |
| 56 | JNK | *Mapk8* | 26419 |
| 57 | Proteoglycans | *Acan* | 11595 |
| 58 | IkB-a | *Nfkbi* | 18035 |
| 59 | SOCS | *Socs1* | 12703 |
| 60 | FOXO1 | *Foxo1* | 56458 |

**Data S1. (separate file)**

Description of the GEO microarray dataset used in the present study. It contains the datasets GSE number with details such as the type of platform, authors, date of first publication, the list of samples, their biological descriptions as well as the binary categorization as OA or WT sample made for the purpose of this study. It also contain a table with the list of mouse genes of potential interest for the model of the current study, with an indication of whether they were present in the final merged dataset (index of gene location in merged dataset if present, NA if absent). The final expression profiles used in the current study contains, as variables, those genes for which an index was found, and the samples from the GEO dataset, as observations.

**Data S2. (separate file)**

Complete profile of computed attractors. 3 attractors were identified for the system of equations described in Supplementary text and computed with the algorithm described in **Fig. S3**. For each attractor, the protein activation level (fast variable), the gene expression level (slow variable) and the global activity (product of the two previous) are indicated for all components of the model.

**Data S3. (separate file)**

Predictions data from the *in silico* screening. All possible pairwise perturbations (i.e. activation or inhibition) were imposed to the hypertrophic state and likehood of transition was evaluated. All conditions that led to a percentage of transition higher than a threshold (70% here) towards the healthy state are reported in this data file. The README sheet provide basic information on the data file content. Predicted conditions are reported in different sheets: the ones reaching the healthy state in 100% of the time, in 99% to 90% of the time, in 89-80% of the time and in 79% to 70% of the time. The column 'Node1' and 'Node 2' indicate the variable index as defined in **Table S2,** the corresponding variable names are indicated in the column 'Names' in the following format: Node1-Node2. The column init_attractor recalls the initial attractor before perturbation, which is the Runx2+ state here, and the column init_value_1 (resp. init_value_2) recalls the initial value in that state for Node1 (resp. Node2). The 'Perturb code' and 'perturbation' columns indicate the type for perturbation for Node1-Node2 as follows: 1 is down-down, 2 is up-up, 3 is down-up, 4 is up-down. The sheet titled 'Summary' report predicted conditions of interest for further experimental validation based on the following criteria: the condition does not involve a transcription factor since their functional activity might be more difficult to target pharmacologically.
